# Population genomics of an emergent tri-species hybrid zone

**DOI:** 10.1101/2022.06.04.494703

**Authors:** Libby Natola, Sampath S. Seneviratne, Darren Irwin

**Author notes:** Corresponding Author: Libby Natola.

## Abstract

Isolating barriers that drive speciation are commonly studied in the context of two-species hybrid zones. There is however evidence that more complex introgressive relationships are common in nature. Here, we use field observations and genomic analysis, including the sequencing and assembly of a novel reference genome, to study an emergent hybrid zone involving two colliding hybrid zones of three woodpecker species: Red-breasted, Red-naped, and Yellow-bellied Sapsuckers (*Sphyrapicus ruber, S. nuchalis*, and *S. varius*). Surveys of the area surrounding Prince George, British Columbia, Canada, show that all three species are sympatric, and Genotyping-by-Sequencing identifies hybrids from each species pair and birds with ancestry from all three species. Observations of mate pair phenotypes and genotypes provide evidence for assortative mating, though there is some heterospecific pairing. Hybridization is more extensive in this tri-species hybrid zone than in two di-species hybrid zones. However, there is no evidence of a hybrid swarm and admixture is constrained to contact zones, so we classify this region as a tension zone and invoke selection against hybrids as a likely mechanism maintaining species boundaries. Analysis of sapsucker age classes does not show disadvantages in hybrid survival to adulthood, so we speculate the selection upholding the tension zone may involve hybrid fecundity. Gene flow among all sapsuckers in di-species hybrid zones suggests introgression likely occurred before the formation of this tri-species hybrid zone, and might result from bridge hybridization, vagrancies, or other three-species interactions.

## Introduction

Speciation is the process wherein an ancestral species subdivides into multiple isolated populations which accumulate differences and eventually become separate species incapable of interbreeding (Coyne & Orr 2004). Understanding how these subdivisions occur and what allows differentiation are fundamental topics in evolutionary biology. Biologists refer to the forces that contribute to differentiation as isolating barriers because they prevent gene flow between the emergent populations. These barriers can be genetic (Dobzhansky 1936), geographic (Mayr 1947), or behavioural (Wells et al. 1978). Contact zones, where differentiating forms come together, provide excellent context for identifying isolating barriers and observing the process of speciation (Barton & Hewitt 1981). The two forms may not interbreed if differentiation is advanced, but early in speciation they might breed and produce offspring (a process called hybridization). In this situation, the contact is called a hybrid zone, and if differentiation between the populations is low, hybrid offspring may breed with each other or backcross with either parental form (a process termed introgression), opening a bridge of gene flow from one species to the other.

Hybridization is typically studied in the context of species pairs (Barton and Hewitt 1985, Moore 1977). However, there are many examples of more complex hybridization relationships in nature (Ottenburghs 2019). In some cases, one species might act as a conduit or a bridge for gene flow between two other species. If the “bridge species” introgresses with both additional species in the “triad”, alleles can be shared between two species indirectly. Bridge hybridization has been observed in suckers (*Catostomus commersoni, C. latipinnis, C. discobolus;* McDonald et al. 2008), *Heliconius* butterflies (*H. melpomene, H. cydno*, and *H. pachinus;* Kronforst 2006) and Darwin’s finches (*Geospiza fortis, G. scandens, G. fulginosa;* Grant and Grant 2020). Similarly, some species may act as introgressive “hubs”, interbreeding with many (otherwise non-interbreeding) species, such as mallard (*Anas platyrhynchos*) hybridizing with many duck species, common pheasant (*Phasianus colchicus*) breeding other Pheasant species, and European herring gull (*Larus argentatus*) which hybridizes with other gulls, potentially facilitating gene flow between pairs of species that do not directly hybridize (Ottenburghs 2019). In other systems, multiple species may hybridize among each other creating more direct introgressive relationships, such as in the *Bos* genus (Wu et al. 2018). Despite there being so many examples of multispecies hybridization, few studies to date have closely examined the phenomenon in bird taxa (Ottenburghs 2019).

Red-breasted, Red-naped, and Yellow-bellied sapsuckers (*Sphyrapicus ruber, nuchalis*, and *varius*) are a recently radiated group of North American woodpeckers. Two well-studied sapsucker hybrid zones exist in British Columbia, Canada: one in the Mackenzie region between Red-breasted and Yellow-bellied Sapsuckers; and another involving Red-breasted and Red-naped Sapsuckers farther south, near the Williams Lake area (Howell 1953, Seneviratne et al. 2016). These zones have historically been separated, but while the Yellow-bellied x Red-breasted zone has been moving south, the Red-breasted x Red-naped zone has crept north. It was long predicted that these two hybrid zones would converge into a single three-species contact zone (Scott et al. 1976, Walters et al. 2002a), but surveys conducted by one of the authors (SS) in the area from 2009-2011 suggested this interaction had not yet occurred.

By 2018, observations of local ornithologists (Dr. Ken Otter 2018, personal communication) and birders (as submitted to eBird; https://ebird.org) suggested the convergence of these hybrid zones had occurred in the Prince George/Quesnel, BC area. Hybrid zones involving three hybridizing species are rare and allow us the opportunity to examine evolutionary dynamics of multiple stages of speciation, often referred to as the speciation continuum (Chhatre et al. 2018, Tarroso et al. 2014). The two colliding zones provide an opportunity to compare hybrid zone dynamics and genomic patterns in the three-species hybrid zone as opposed to the two paired species zones. This allows us to directly differentiate patterns of three species gene flow from two.

In this study, we use field and molecular methods to describe multispecies hybridization among sapsuckers. We aim to answer the questions: 1) to what degree do Red-naped, Red-breasted, and Yellow-bellied Sapsuckers hybridize in sympatry? and 2) how do these interactions vary among different hybrid zones? We address these questions with surveying, mate pair, and genomic data of sapsuckers in the putative tri-species hybrid zone. We compare patterns of introgression in this zone with those in each of the two-species hybrid zones.

## Materials and Methods

### Surveying

To assess sapsucker density and habitat preferences we performed standardized presence/absence surveys between 5 and 10 AM during the months of May and June in 2019 and 2020. We did not survey in rainy, cold, or windy weather. We surveyed residential and Forest Service roads by stopping once every kilometer and playing a standard recording on a Bluetooth speaker. This recording included two minutes of silence, 3.5 minutes of drumming and call noises from Red-breasted, Red-naped, and Yellow-bellied sapsuckers, and two more minutes of silence. During the silences, we listened for sapsucker calls and drumming, and looked for sapsuckers. We marked each survey point as either having sapsuckers present or absent. If sapsuckers were present, we identified them to species based on plumage markings. Hybrids could not be identified to species cross because Red-breasted x Red-naped hybrids and Red-breasted x Yellow-bellied hybrids cannot be reliably differentiated by plumage. We identified mated pairs as those seen excavating, feeding, or defending a nest together, or responding to call playback with paired dry chatter and moth flight display (Walters et al. 2002b).

### Sample collection

To reflect the population-wide genotypic composition in the tri-species hybrid zone, we attempted to catch every sapsucker we found, regardless of phenotype, and made a concerted effort to fill in any geographical sampling gaps. We trapped 107 birds along two intersecting transects, one ~115 km N-S on highway 97 from north of Bear Lake, BC to just south of Soda Creek, BC and another spanning ~140 km E-W between Vanderhoof, BC and McBride, BC on highway 16. We caught sapsuckers using either dipnets at a nest or canopy nets accompanied by audio and decoy lures. In hand, we collected ~ 50μL of blood from the brachial vein of each bird and stored it in Queen’s lysis buffer. We banded each individual for later observation and to prevent resampling. We also collected morphometric data and photographed each bird. Woodpeckers replace feathers in a reliable sequence as they age, so we determined age classes based on wear and replacement in the secondary and primary covert feathers (Pyle 1997, pp 163–181). For analyses, we collapsed age classes to second year (SY, bird hatched previous summer) or after second year (ASY, bird hatched prior to previous summer). In this study we analyze our tri-species hybrid zone data along with data from two di-species hybrid zones sampled by Seneviratne et al. (2016, 2012): one Red-breasted x Yellow-bellied Sapsucker zone and one Red-breasted x Red-naped Sapsucker zone. For sample collection, DNA extraction, and GBS sequencing methods of di-species hybrid zones please reference Seneviratne et al. (2016, 2012).

### Reference genome assembly

To create a *de novo* reference genome, we submitted tissues of one female Red-breasted Sapsucker voucher specimen to the University of Delaware Sequencing & Genotyping Center for Pacific Biosciences Single Molecule Real Time (SMRT) Consensus Long Read (CLR) sequencing on a Pacific Biosciences Sequel II sequencer. Sequencing yielded 172X coverage. We assembled the genome in canu v. 1.9 (Koren et al. 2017), resulting in a 1.6 Gb assembly with an N50 of 4.7 Mb, and an LG50 of 75, with 7596 contigs. To make the assembly haploid for downstream analysis we kept one haplotig from each of the heterozygous contigs with the Purge Haplotigs pipeline (Roach et al. 2018). We then masked repetitive elements using Repeat Masker v. 4.1.1 (Smit et al. 2013), which marked 15.93% of the genome as repeats, the majority of which (13.71% of the genome) are LINE retroelements. We assessed completeness of our assembly with the program BUSCO v. 5.1.2 (Manni et al. 2021) and the aves_odb10 lineage dataset with 8338 BUSCOs. Our assembly is 96% complete with 88% complete single copy BUSCOs, 8% complete duplicated, 0.9% fragmented, and 3.1% missing. Finally, we used the Chromosemble tool in Satsuma v. 2.0 (Grabherr et al. 2010) to align our contigs to a Golden-fronted Woodpecker (*Melanerpes aurifrons*) genome (Wiley & Miller 2020). Our final assembled genome had 269 scaffolds.

### GBS protocol

We extracted DNA using a standard phenol-chloroform extraction and subsequently followed the genotyping-by-sequencing (GBS) protocol detailed by Elshire et al. (2011) and Alcaide et al. (2014) using the enzyme *PstI*. We followed lab protocols of Seneviratne et al. (2016) with the following changes. We incubated the digestion with 1 μL *PstI*, 2 μL 10X buffer, 6μL of common adaptor, 6μL of barcode, and 5 μL of 20ng/μL template DNA at 37°C for 2 hours. We ran the ligation incubation at 22°C for 1 hour then 65°C for 10 minutes. For PCR, we included 5μL of 5x Phusion buffer, 0.5 μL 10mM dNTPs, 1.25 μL forward and reverse primers, 12.75 μL UltraPure water, and 0.25 μL of PhusionTaq. We ran the reaction protocol as follows: 98 °C for 30 seconds and 18 cycles of: 98°C for 10sec, 65°C for 30sec and 72°C for 30 sec. This is followed by an extension of 72°C for 5min, followed by 4°C. We used gel extraction to select for 400-500 bp fragments. We sent 100 ng of DNA from 289 sapsuckers (Supplemental Materials Table 1) to Genome Quebec for sequencing on either an Illumina HiSeq4000 PE150 or the NovaSeqSP 6000 PE150 (Table S1) (Illumina, San Diego, CA).

### GBS Processing

We demultiplexed all reads using a custom script from Irwin et al. (2018), trimmed reads with Trimmomatic v 0.38 (Bolger et al. 2014), then aligned them to our Red-breasted Sapsucker reference genome using BWA v 0.7.17 (Li 2013). We further realigned around indels and called genotypes using GATK v 3.8 (Van der Auwera & O’Connor 2020). We kept variant sites and removed all indels and sites that had mapping quality < 20, heterozygosity > 0.6, more than 70% missing data, GQ < 10, or minor allele frequency ≥ 0.05. We calculated φ in VCFtools v 0.1.16 (Danecek et al. 2011) to estimate relatedness, randomly removed one individual from each pair of birds with an estimated 4^th^ degree relationship or closer (phi ≥ 0.0224), and filtered out individuals with more than 60% missing data. After filtering, we retained 251 individuals and 50,639 SNPs.

### Data analysis

Using the variants data, we visualized genomic relationships by generating a genomic PCA in R (R Core Team 2011) using scripts adapted from Irwin et al. (2018). To generate ADMIXTURE plots, we further pruned data for linkage disequilibrium, removing sites with > 0.6 correlation in 1000 bp windows. We ran ADMIXTURE v 1.3.0 (Alexander & Lange 2011) for *K* = 1-6 with cross validation and calculated standard errors using 2000 bootstrap replicates. Cross validation error indicated *K* = 3 had the best predictive accuracy, consistent with the expectation of this being a three-species hybrid zone. We used the R package Ternary v 1.2.2 (Smith 2017) to create ternary plots of ADMIXTURE data.

### Assortative Mating

We evaluated the strength of assortative mating (AM) among the three species in the tri-species hybrid zone with plumage-based species assignment of birds from 91 sighted pairs. We classified all birds to pure species or hybrid by plumage, and because different hybrid species crosses are indistinguishable by eye all hybrids are lumped. First, we calculated the proportion of conspecific pairs we found in the population (including pairs involving two phenotypic hybrids). To evaluate whether this proportion falls within expectations under random mating, we randomized pairings based on observed species and hybrid frequencies 1000 times and calculated the proportion of conspecific pairs for each randomization. We checked if the conspecific proportion in our data was within the upper 5% of the simulated distribution. Then, we calculated plumage AM using the equation:

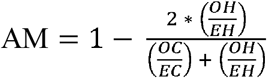

where *OH* signifies observed heterospecific pairings, *EH* is expected heterospecific pairings, *OC* is observed conspecific pairings, and *EC* is expected conspecific pairings, in accordance with methods in Scordato et al. (2020). We were able to capture and genotype both mates from 15 pairs. We charted the ADMIXTURE K=3 cluster data of the 15 genotype pairs on a Ternary plot and measured the Euclidian distances between each pair’s Cartesian coordinates on the Ternary plot. For each individual, we plotted its p coefficient (proportion of genome attributed to Red-breasted Sapsucker ancestry in ADMIXTURE) by its measured genotypic mate distance. We ran a linear regression to model predictability of genotypic mate distance based on p coefficient. Our reasoning with this latter analysis is that the majority of the birds in the hybrid zone are phenotypic Red-breasted Sapsuckers, such that they may on average mate with more similar individuals than the other phenotypes are able to.

### Hybrid Fitness

To better understand hybrid performance in the tri-species hybrid zone, we asked whether there is evidence that degree of ancestry admixture is associated with survival. We did this by examining whether different age classes differed in their average amount of admixture. We were unable to collect hatching or nestling data in the field, so we first predicted the ADMIXTURE cluster genotypes of birds in the next generation from our genotypic mate pair data to see if hybrid sapsuckers were equally represented after one year as we might expect from breeding data. We averaged ADMIXTURE clusters p, q, and z among mated pairs to predict offspring ancestry, then calculated a measure *m*, calculated as 1 minus the maximum value across all three ADMIXTURE clusters, for these “hatch year” birds. Next we distilled admixture within all sequenced birds to *m* and plotted this measure against sapsucker age class (HY, SY, or ASY) to understand if hybrids are less likely to survive to older ages, as might be predicted if post-zygotic isolation causes them to be less fit than pure birds. In addition, we ran a logistic regression on genomic admixture (*m*) and age class to see if amount of admixture predicted likelihood of surviving to older age classes.

## Results

The genomic data support the hypothesis that there is a three-species hybrid zone in east-central British Columbia. In the PCA (Fig. 2), each species formed a distinct group with the allopatric samples at the distal ends of each cluster. The tri-species hybrid zone samples showed each species formed a separate genomic cluster. The main axis of variation (PC1, 16.3%) identifies differences between Yellow-bellied Sapsuckers and Red-breasted/Red-naped Sapsuckers, and these latter sister species split out across PC2 (3.9%) (Fig. 2). It is important to note that the relative density of each species varies widely. There is a much higher proportion of phenotypically Red-breasted birds (70.4% surveyed, 53.5% sampled) than any other species, followed by phenotypically hybrid birds (20.2% surveyed, 29.1% sampled). We found relatively few Red-naped (4.0% surveyed, 8.1% sampled) and Yellow-bellied (5.4% surveyed, 9.3% sampled) birds in the hybrid zone.

**Figure 1.**
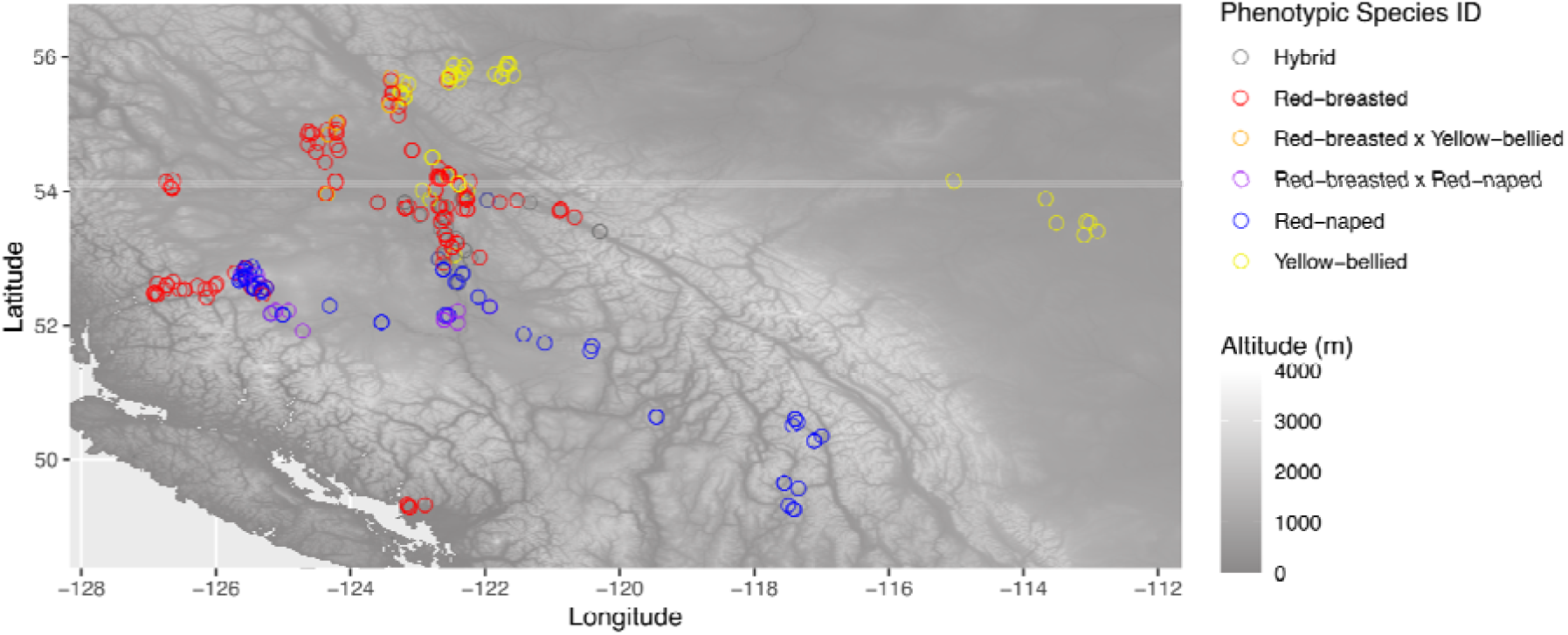
Map of sample locations for phenotypic hybrids (gray) Red-breasted (red), Red-breasted x Yellow-bellied (orange), Red-breasted x Red-naped (purple), Red-naped (blue), and Yellow-bellied (yellow) sapsuckers across southeast Canada.

**Figure 2.**
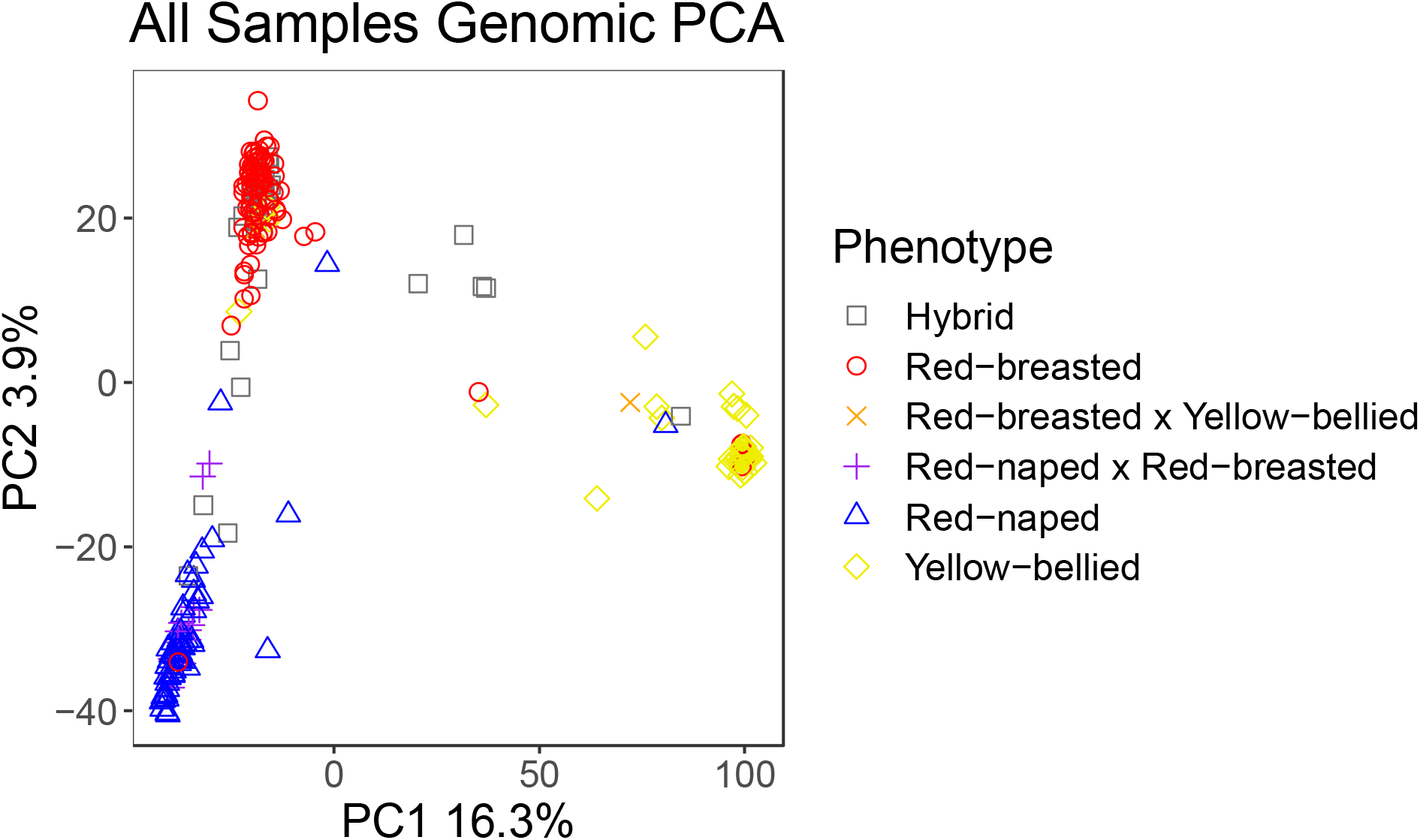
Principal Component Analysis of genomic data, each sample is depicted by its phenotypic species assignment including tri-species hybrid zone hybrids (grey squares), Red-breasted (red circles), Red-naped (blue triangles), Yellow-bellied (yellow diamonds), hybrids from Re-breasted x Yellow-bellied di-species hybrid zone (orange x), and hybrids from Red-naped x Red-breasted di-species hybrid zone (purple +).

There are genomically intermediate birds placed between each pair of species clusters, with the most being between Red-breasted and Red-naped, and the fewest intermediates between Red-naped and Yellow-bellied (Fig. 2). Intermediate Red-breasted/Red-naped birds fell along a wide gradient of PC2 values, but there was a distinct gap separating the two species’ clusters. Intermediates involving Yellow-bellied sapsuckers tended to be more clumped in the very center of the X-axis, equidistant from each parent cluster. The majority of birds placed between the major species clusters were collected in the tri-species hybrid zone. Finally, though most birds were placed in genotypic clusters that matched their phenotypic assignment, there were genotypic/phenotypic mismatches, shown in the PCA as samples that are the “wrong” color for their predicted species cluster, pure looking birds with hybrid genotypes, or intermediate looking birds with pure genotypes. Most mismatched birds were from the tri-species hybrid zone.

ADMIXTURE results similarly showed that many individuals in the study belong to one of the three distinct species (clusters of ancestry values q, p, or z > 0.995, 14 Red-breasted, 21 Red-naped, 38 Yellow-bellied) (Fig. 3). There were also many birds with both Red-breasted and Red-naped ancestry (q + p > 0.995, 57 birds). Several birds showed primarily Red-breasted x Yellow-bellied ancestry (q + z > 0.995, 10 birds), and seven had primarily Red-naped x Yellow-bellied ancestry (p + z > 0.995). The biggest group of birds had a combination of all three ancestries (116 birds, hereafter termed “muttsuckers” after McDonald et al. (2008)), many of which had a low proportion of Yellow-bellied, some Red-naped, and a high proportion of Red-breasted ancestry. In this way the ADMIXTURE explicitly shows the three-species ancestry found in many birds in this region.

**Figure 3.**
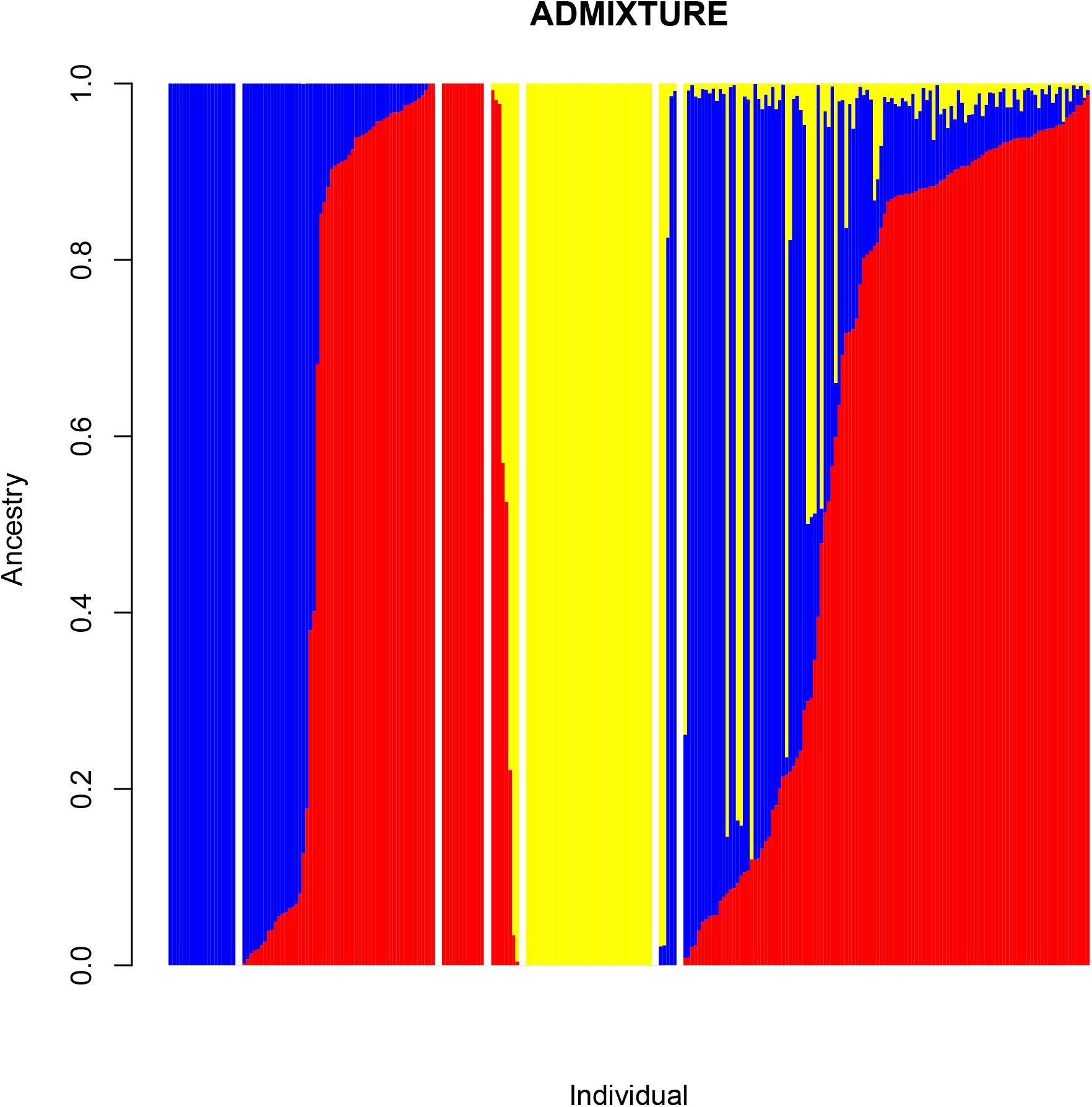
Plot of ADMIXTURE k = 3 data for all samples. Ancestry coefficient was partitioned to Red-breasted (q, red), Red-naped (p, blue), and Yellow-bellied (z, yellow) clusters. Plot is broken up to show pure birds from each cluster, each species pair cluster, and birds with ancestry from all three species.

The ternary plot shows these data more intuitively, with many birds placed on the axis shared by Red-breasted and Red-naped, especially towards the Red-breasted corner, a few along the Yellow-bellied axes, and many points within the triangle, signifying three-species ancestry (Fig. 4). With the tri-species hybrid zone, di-species hybrid zones, and allopatric zones birds split into separate Ternary plots, it is clearer that the three species are well isolated in allopatry (Fig. 4a), and there is increasingly more admixture as we move from di-species hybridization (Fig. 4b) to tri-species hybridization (Fig. 4c). It’s also more obvious that we caught no “pure” Red-naped Sapsuckers in the tri-species hybrid zone. Again, there are fewer intermediates showing recent Yellow-bellied ancestry, and those that do exist tend to be equidistant from either parental group. The ternary plot also reiterates the “mismatch” pattern in which phenotypic assignment does not consistently mirror genotypic assignment.

**Figure 4.**
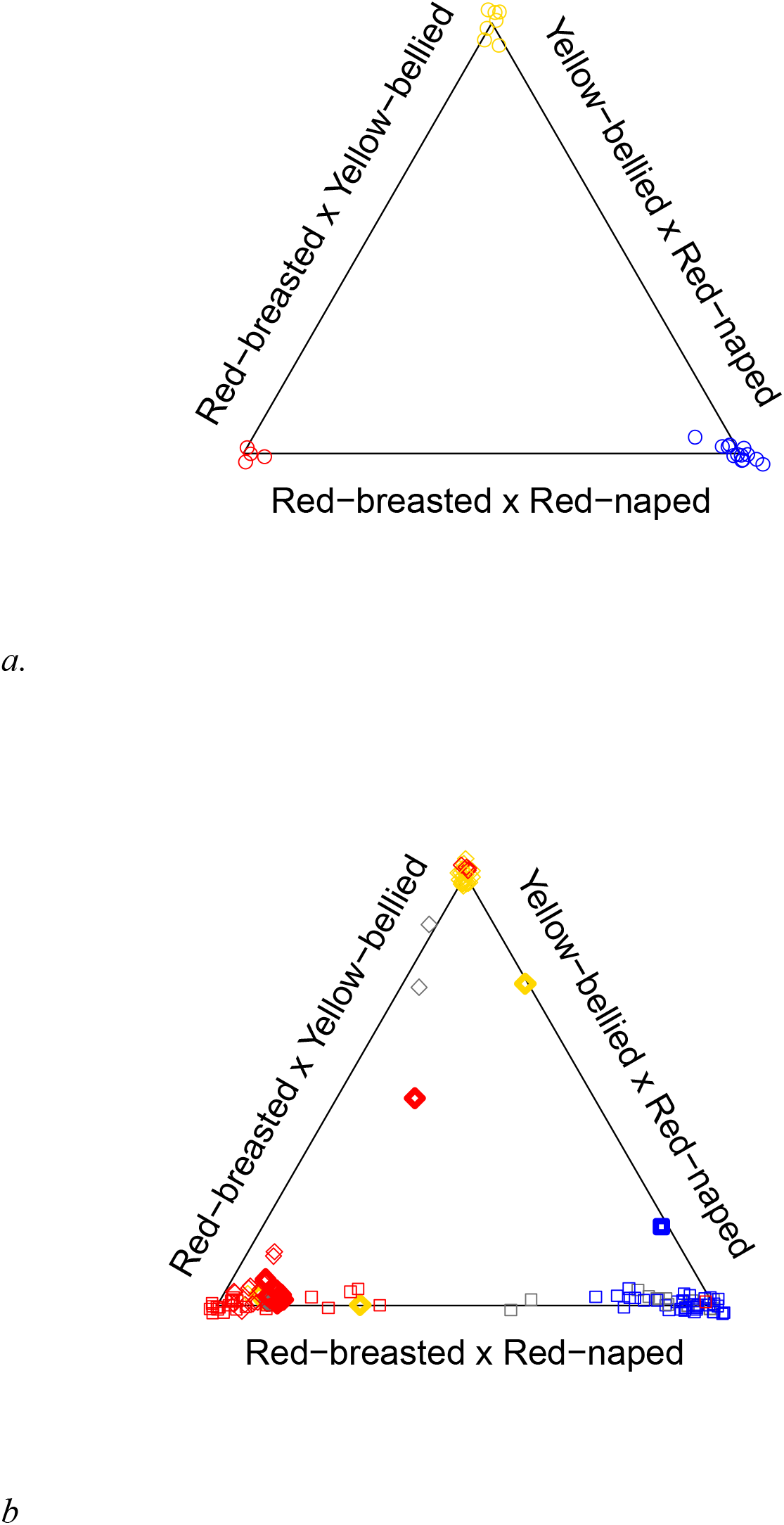
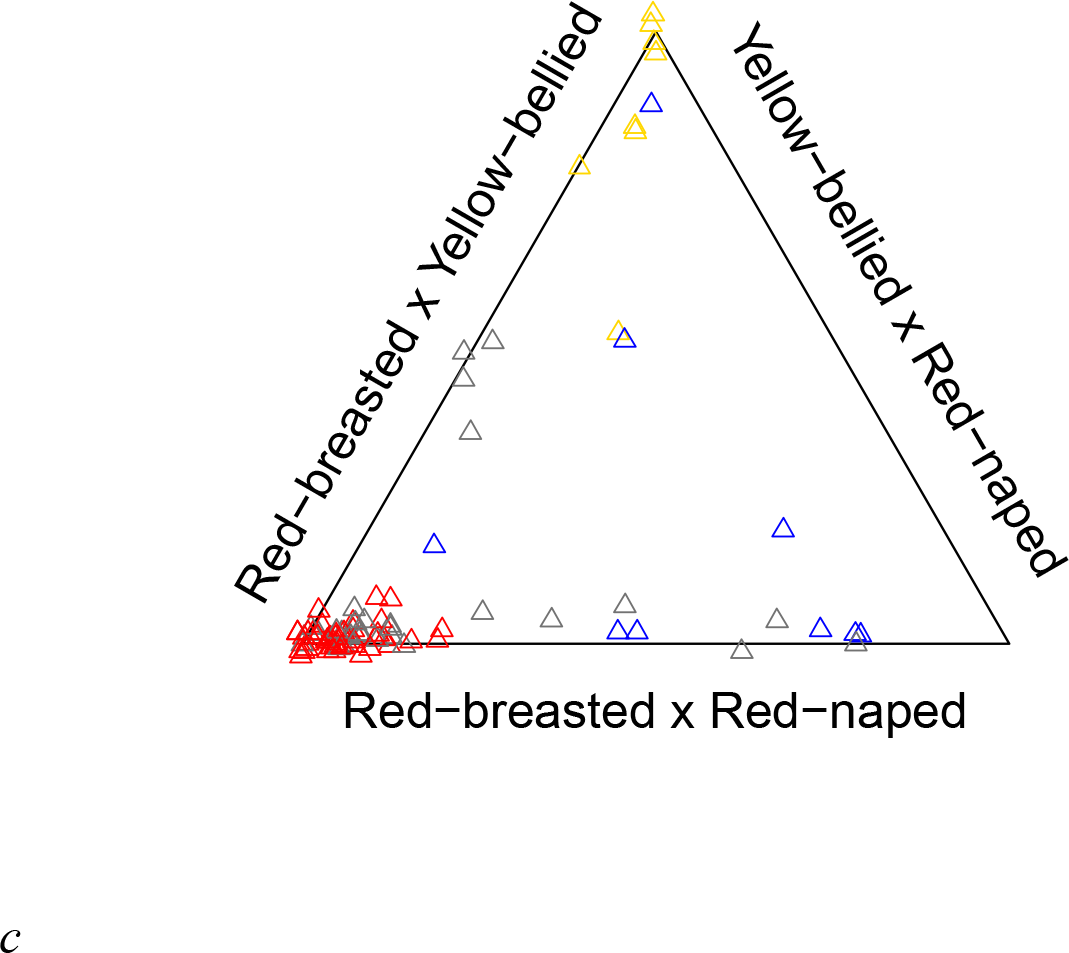
Ternary plots for allopatric zone (a), di-species hybrid zones (b), and tri-species hybrid zone (c) samples. Sample colors depict phenotypic species assignment for Red-breasted (red), Red-naped (blue), Yellow-bellied (yellow), and hybrid (grey) birds. In panel b, the Red-breasted x Red-naped di-species hybrid zone samples are shown as squares and the Red-breasted x Yellow-bellied di-species hybrid zones are shown as diamonds, and birds with unexpected genotypes given their sampling location are shown in bold. Points are jittered to show all samples.

Based on the observed species frequencies, the expected proportion of conspecific pairings is 0.56. Our pairing randomization showed the observed proportion of conspecific pairings (0.67) falls in the 99.7th quantile of our randomized pairing distribution, which gives us high confidence that sapsuckers are mating conspecifically more frequently than would be expected under random mating (p <0.01). The AM calculations show that matings between phenotypic species are under-represented, but matings between “pure” individuals and plumage intermediates are more common (Table 1). The genotype mate pairs on the ternary plot have short genomic distances between mate pairs near Red-breasted Sapsuckers, and much longer genomic distances connecting pairs with hybrids and lower-density (Fig. 5a). The scatter plot of p coefficient of randomly chosen mate by pair’s genomic distance (Fig. 5b) shows a strong negative correlation (R^2^ = 0.8, F = 52.1, *p* < 0.0001), meaning that the more Red-breasted ancestry a bird has, the more closely related it tends to be to its mate.

**Table 1.**
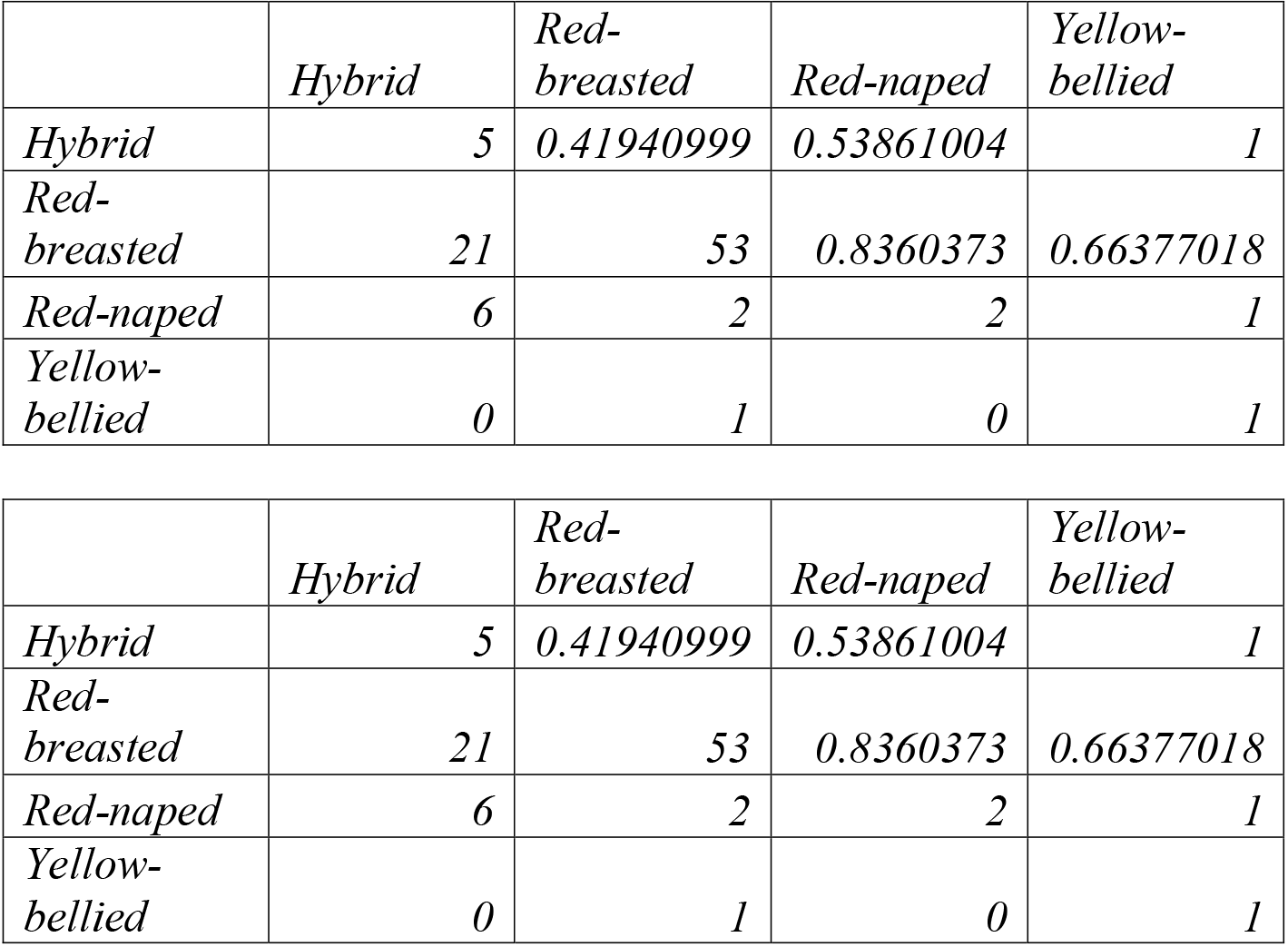
Assortative mating matrix based on phenotypic data. Values on and below the diagonal represent the number of pairs found for each cross type. Above the diagonal, italicized, are the calculated AM values indicating the degree of assortative mating between each pair from random mating (0) to complete isolation (1) or completely disassortative mating (−1).

**Figure 5.**
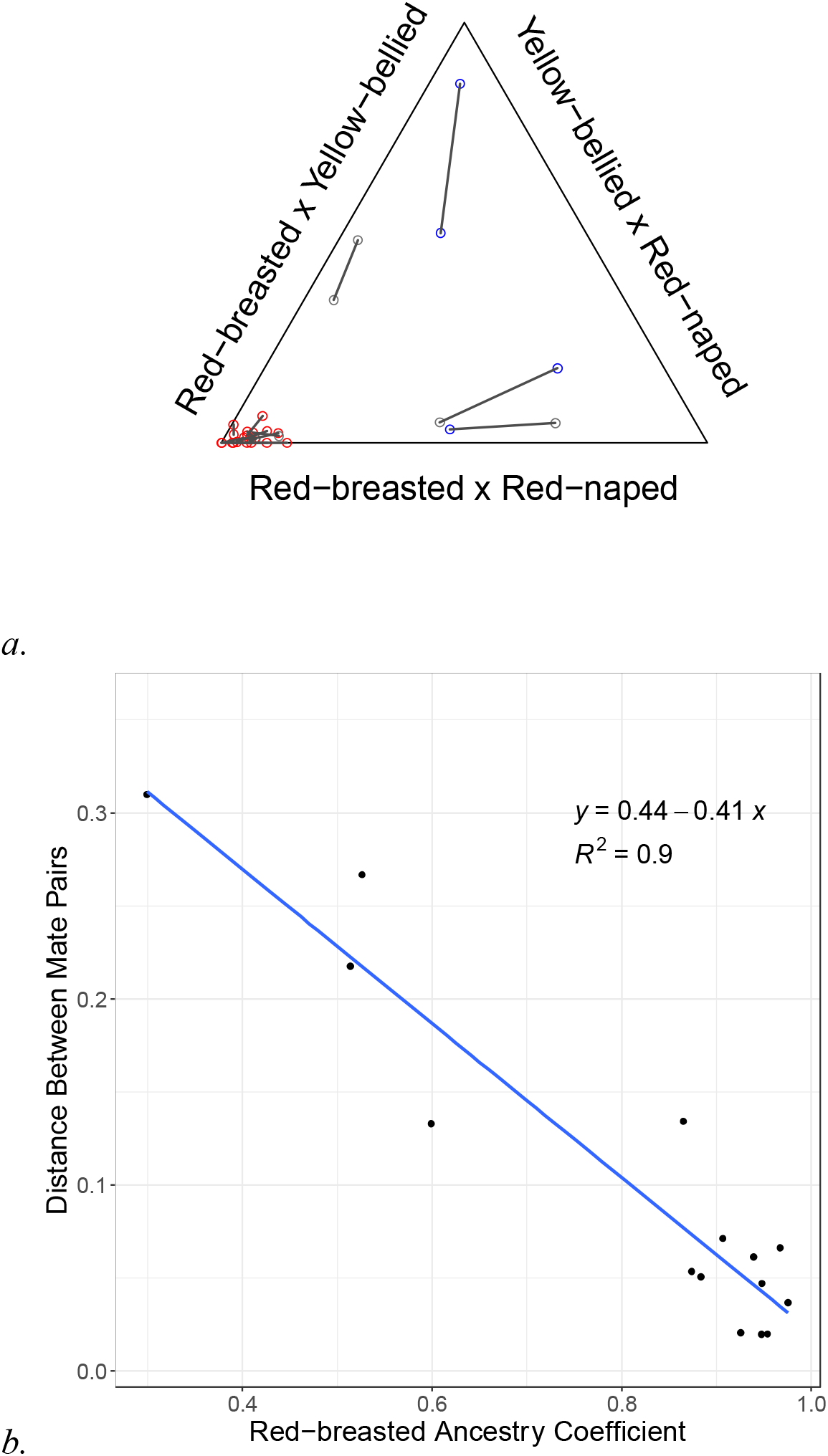
Ternary plot of mate pair genotypes from tri-species hybrid zone (a). Colors depict phenotypic species assignments for Red-breasted (red), Red-naped (blue) and hybrid (grey). Grey lines connect paired mates. Proportion of Red-breasted Sapsucker ancestry by Euclidian distance between mate pairs’ Cartesian coordinates (b) shows one mate chosen randomly per pair along with linear regression line, equation, and R^2^ value. Shaded areas indicate 95% confidence level interval.

Looking at age classes as predicted by genomic admixture, we found that hybrids were not over-represented in the SY or ASY age class (HY *t* = 0.11, *p* = 0.91, SY *t* = −0.85, *p* = 0.4, Fig. 6), so we have no evidence hybrids suffer lower survival rates than pure species.

**Figure 6.**
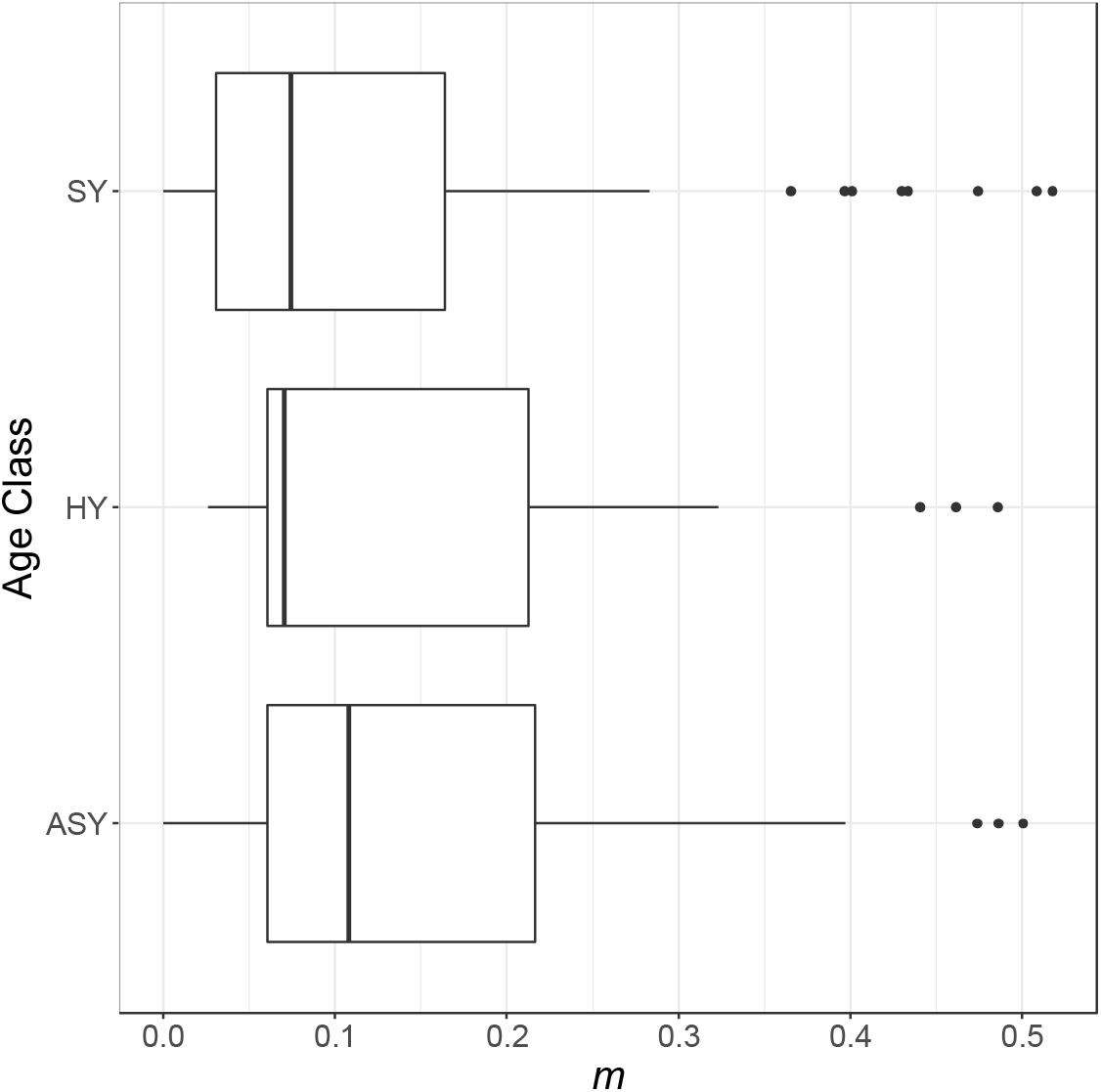
Box and whisker plots showing measured admixture (m), for projected hatch year (HY) birds, and plumage identified second year (SY) and after second year (ASY) birds.

## Discussion

### Di-vs Tri-species Hybrid Zone Dynamics

Our survey and genomic data conclusively show there is a tri-species hybrid zone in the Prince George region of BC. Individuals from all three species breed here, often near one or both of these other sapsucker species. However, we note that the great majority of Red-naped sapsuckers in the tri-species hybrid zone have some admixture. There is also a demographic imbalance, with high representation of Red-breasted phenotypes and genotypes. We present evidence that the sapsuckers breed in all three species pairs in the tri-species hybrid zone, and there appears to be back-crossing in every direction, indicating direct gene flow and introgression among all three species. Additionally, we found many “muttsuckers” with ancestry from all three species, meaning there is likely also indirect gene flow as well, in which some geneflow could occur between two species that does not involve them directly interbreeding if hybrids of two species breed with a third species. Backcrossed admixed birds (largest ADMIXTURE cluster < 0.76) account for 18.9% of birds sampled in the tri-species hybrid zone, and non-backcrossed hybrids account for 3.3% (largest cluster < 0.51). However, allopatric individuals from each species form distinct groups (Fig. 4a), showing that the evidence for substantial introgression is limited to the sympatric region.

Compared to the tri-species hybrid zones, we see less hybridization and introgression in general within the di-species hybrid zones (as in Grossen et al. 2016, Seneviratne et al. 2016). There are very few that could be direct hybrids (0.7% had a maximum ADMIXTURE cluster < 0.51), and less than 7% are recently admixed (largest cluster < 0.76), showing that the evidence for substantial introgression is limited to the sympatric region. As expected by their closer evolutionary relationship, Red-breasted and Red-naped hybridize the most, with mitochondrial genetic distance producing estimates of divergence times of 1.090 million years ago (Yellow-bellied) and 0.32 million years ago (Red-breasted and Red-naped) (Weir & Schluter 2004). Very little introgression seems to occur between either species and Yellow-bellied, but it is important to note that there are early generation backcrosses between Yellow-bellied and both Red-naped and Red-breasted, so at least some hybrids can be viable and fertile. It is intriguing that there is more admixture in the tri-species hybrid zone than the di-species hybrid zones. Our data don’t have the power to explain this pattern, so it opens some interesting new questions. Does the addition of a third species reduce assortative mating? Could the distorted demographics in the tri-species hybrid zone affect mate availability in low-density species, therefore promoting hybridization? We know habitat influences hybrid zone dynamics in sapsuckers (Natola et al., *in preparation).* The large stretch of aspen dominated valleys of Prince George area is surrounded by conifer dominated hills. The latter is less frequented by sapsuckers (Sampath Seneviratne, *personal observation*). Perhaps climate or landscape features in this region inherently promote hybridization.

Interestingly, in the di-species hybrid zones there are several individuals that have ancestry from the third species that is not one of the two species apparently hybridizing there. For instance, there are 13 samples from the Red-breasted x Yellow-bellied hybrid zone (bolded diamonds on Figure 4b) with more Red-naped ancestry than Yellow-bellied ancestry, in addition to a bird from the same hybrid zone with Red-naped but no Red-breasted ancestry, and a bird from the Red-breasted x Red-naped zone (bolded square Fig. 4b) with no Red-breasted ancestry but appears to be a relatively recent Red-naped x Yellow-bellied backcross. Also, one muttsucker from the Red-breasted x Yellow-bellied hybrid zone (bolded diamond Fig.4b) has substantial ancestry from all three species. The existence of genotypes in the di-species hybrid zones in which they don’t “belong”, suggests greater introgression than previously reported. These samples were all collected approximately 10 years before local ornithologists noticed the convergence of the two zones. In addition, we see evidence of many advanced generation muttsuckers (at least one species’ ancestry > 0%, but < 10%). With an estimated generation time of 1.9 years (Seneviratne et al. 2016), we expect the eight years, or fewer than 5 generations, since sampling showed no tri-species contact, is insufficient to produce the proportion of late generation backcrosses with tri-species ancestry we found. It may be possible that the Red-breasted Sapsuckers, which have existed in allopatry between the two hybrid zones, have acted as a conduit for introgression of alleles between all three species for generations. Alternatively, as two di-species hybrid zone birds appear to have very little Red-breasted ancestry at all, and a Red-breasted x Red-naped hybrid from the Red-breasted x Yellow-bellied hybrid zone has very little Yellow-bellied ancestry (Fig. 4b), there may be intermittent contact through occasional long-distance dispersal. In any case, the data showing existence of muttsuckers or birds with ancestry of species excluded from their hybrid zone is evidence that three-way gene flow among all three species predates our detection of the tri-species hybrid zone.

### Tri-species hybrid zone selection

Though there is a lot of hybridization and introgression in the tri-species hybrid zone, and to a more limited extent throughout the di-species hybrid zones, there is no evidence of a collapse into a hybrid swarm. If birds were mating randomly and hybrids were as fit on average as the parental species, the ternary plot (Fig. 4c) would eventually show most birds in the middle and few at the corners. Instead, our samples are overwhelmingly concentrated to the corners and edges of the plot. We propose that the Prince George, BC region is therefore a three-way tension zone (Barton & Hewitt 1989), maintained by reduced migration in and out of the hybrid zone and selection against hybrids or hybridization itself. Therefore, we suspect some form of selection against hybrids, perhaps enhanced by assortative mating, is maintaining species boundaries and reducing gene flow in this system.

We investigated evidence for assortative mating. In regard to plumage, sapsuckers are not mating randomly, as evidenced by the higher proportion of conspecific pairings than random, and the high AM values above the diagonal in Table 1, suggesting some amount of mate preference for phenotypically similar individuals (note, however, low sample sizes of Yellow-bellied Sapsuckers in the AM analysis). This provides evidence for a partial mating barrier between the species. That holds up with the genomic data, where we saw few F1 hybrids (Fig. 4c). However, there appears to be little isolation between phenotypic hybrids and phenotypic species. Assortative mating was high with Yellow-bellied Sapsuckers, but we had a very small sample size for this species in these analyses. Assortative mating rates were far lower between hybrids and pure Red-breasted or hybrids and pure Red-naped Sapsuckers. This is supported by our evidence of several backcrosses given the number of F1s we found, that hybrids may be acceptable mates. Note however, the apparent offset of plumage phenotype and genotype in many birds. Given this offset and acceptance of hybrid birds, it is likely that sapsucker mate choice is not based solely on plumage. These patterns also suggest plumage may be controlled by a few genes of large effect, making it an excellent candidate for a genome-wide association study.

The analysis of the 15 genotyped pairs shows that individuals with Red-breasted genotypes tend to pair with each other, and hybrids and Red-naped genotypes (no Yellow-bellied were in this analysis) tend to mate with far more genetically distant sapsuckers (Fig. 5a, b). These patterns could be explained by a preference for genetically similar individuals. We speculate this is an effect of relative demography. There are so few Red-naped in the area that those who are looking for mates “settle” for a mate with an ill matched genome rather than foregoing breeding entirely. However, with as much outcrossing as we see in the full genomic dataset it is unlikely assortative mating alone can explain the overall species maintenance (Irwin 2020), so we speculate there must be some selection against hybrids as well. Sources of low hybrid fitness against hybrids might be related to habitat specialization, as the different sapsucker species have different preferred climatic conditions and forest compositions and hybrids could be at a disadvantage (Natola & Burg 2018, Walters et al. 2002b, 2002a). Seasonally migratory behaviour might also cause low hybrid fitness, as all three of these species have different migratory strategies (Walters et al. 2002b, 2002a) and intermediate routes could be disadvantageous (Bensch et al. 1999, Delmore et al. 2016, Scordato et al. 2020). Additionally, there could be genomic incompatibilities which decrease hybrid fitness.

We investigated one component of hybrid fitness, survival from hatching to adulthood, by examining age classes and admixture rates. We didn’t find any evidence that hybrids are less likely to survive to adulthood based on our mate pair data, or that second year birds are less likely to survive to after second year based on our plumage data (Fig. 6). The n = 15 for our mate pair data is small and may have affected our ability to detect true patterns. While we didn’t find evidence of reduced hybrid survival, it is possible hybrids produce fewer offspring on average and their fitness is thus diminished. Genetic incompatibilities, caused for instance by copy number variation and chromosomal inversions, could explain a decrease in hybrid fecundity as these interactions typically affect the zygotes formed from heterokaryotypic gametes (Rieseberg & Willis 2007). More research into hybrid fecundity and genomic synteny among sapsucker species could clarify whether these interactions may be lowering fitness of hybrids.

### Conclusions

Our data show that increasing the number of hybridizing species from 2 to 3 changes hybridization dynamics. Hybridization and introgression are more pervasive in the tri-species hybrid zone than the di-species hybrid zones. However, that introgression does not extend to allopatric populations, and hybridization is less common than expected under random mating. Therefore, this region serves as a tri-species tension zone. As yet, it is unclear what sources of selection against hybrids maintains the three species in the hybrid zone, yet hybrid survival to adulthood doesn’t seem to explain the pattern. A genomic study on loci under selection and resistant to introgression may clarify the underlying causes of isolation in sapsuckers. Furthermore, we show that hybridization is more complex than traditional two-species interactions in sapsuckers. Indirect gene flow between the three species appears to predate the tri-species hybrid zone we identified, meaning this history of triple introgression might be the result of conduits, vagrancies, or other three-species contact. This signifies that sapsucker species interactions were more complex than we previously suspected, and may indicate a more pervasive biological pattern present in other taxa.

## Acknowledgements

Kathy Martin, Loren Rieseberg, and Dolph Schluter provided key feedback on all stages of project development. Afnan Ali, Avery Bartels, Jamie Clarke, Gabriel Pang, Rashika Ranasinghe, and Kang Wang assisted in field sample collection. This research was funded by the Natural Sciences and Engineering Research Council of Canada (RGPIN-2017-03919 and RGPAS-2017-507830 to DI), the Society of Canadian Ornithologists’ Baillie Award, the American Museum of Natural History’s Chapman Award, the Explorer’s Club’s Mamont Grant, the Society for the Study of Evolution’s R.C. Lewontin Early Award, and the Warner and Hildegard Hesse Research Award. Thanks to Gavin Hanke (Royal BC Museum) Jocelyn Hudon (Royal Alberta Museum), Chris Stinson and Ildiko Szabo (Beaty Biodiversity Museum), and Kevin Winker (University of Alaska Museum) for sharing tissue samples.

## Data Accessibility

All scripts, data, and analyses are available on a GitHub repository, and genetic data will be posted to NCBI. Both will be made public upon manuscript acceptance.

## Benefits Generated

Benefits from this research accrue from the sharing of our data and results on public databases as described above.

## Author Contributions

LN and DI conceived the project. LN assembled reference genome, sampled and prepared tri-hybrid zone GBS libraries for sequencing, processed reads and conducted analyses, with advice from DI. SS sampled and prepared GBS libraries from both di-hybrid zones. LN wrote the manuscript with input from DI and SS.

