## Supplemental Materials Table 1 for "Population genomics of an emergent tri-species hybrid zone" (from another version of this preprint: version 1, 2022-06-04)

Table S1: Sample ID, phenotypic species assignment, capture date, latitude, longitude, phenotypic sex, sampling zone (tri-species hybrid, di-species hybrid, or allopatric), and sequencer used to generate GBS reads.

| ID | Species | Date | Lat | Lon | Sex | Zone | Sequencer |
| --- | --- | --- | --- | --- | --- | --- | --- |
| PG_2019_TE02L01 | Red-breasted | 2/5/19 | 53.7701 | -123.1331 | u | tri | HiSeq 4000 PE150 |
| PG_2019_TE05L01 | Red-breasted | 5/5/19 | 54.1829 | -122.6636 | u | tri | HiSeq 4000 PE150 |
| PG_2019_TE05L02 | Red-breasted | 5/5/19 | 54.1829 | -122.6636 | u | tri | HiSeq 4000 PE150 |
| PG_2019_TE05L03 | Red-breasted | 5/5/19 | 54.2007 | -122.6949 | u | tri | HiSeq 4000 PE150 |
| PG_2019_TE06L02 | Red-breasted | 5/6/19 | 53.8765 | -122.272 | u | tri | HiSeq 4000 PE150 |
| PG_2019_TE06L03 | Red-breasted | 5/6/19 | 53.8765 | -122.272 | u | tri | HiSeq 4000 PE150 |
| PG_2019_TE06L04 | Red-naped | 5/6/19 | 53.8697 | -121.9672 | f | tri | HiSeq 4000 PE150 |
| PG_2019_TE07L01 | Yellow-bellied | 5/7/19 | 53.77 | -122.6341 | m | tri | HiSeq 4000 PE150 |
| PG_2019_TE07L02 | Hybrid | 5/7/19 | 53.77 | -122.6341 | u | tri | HiSeq 4000 PE150 |
| PG_2019_TE07L03 | Hybrid | 5/7/19 | 53.77 | -122.6341 | u | tri | HiSeq 4000 PE150 |
| PG_2019_TE07L04 | Red-breasted | 5/7/19 | 53.7645 | -122.5751 | u | tri | HiSeq 4000 PE150 |
| PG_2019_TE08L01 | Red-breasted | 5/8/19 | 53.894 | -122.3194 | u | tri | HiSeq 4000 PE150 |
| PG_2019_TE08L02 | Red-breasted | 5/8/19 | 53.894 | -122.3194 | u | tri | HiSeq 4000 PE150 |
| PG_2019_TE08L03 | Hybrid | 5/8/19 | 53.894 | -122.3194 | u | tri | HiSeq 4000 PE150 |
| PG_2019_TE11L01 | Hybrid | 5/11/19 | 53.0021 | -122.2234 | u | tri | HiSeq 4000 PE150 |
| PG_2019_TE11L02 | Red-naped | 5/11/19 | 52.9948 | -122.69 | f | tri | HiSeq 4000 PE150 |
| PG_2019_TE11L03 | Hybrid | 5/11/19 | 52.9948 | -122.69 | u | tri | HiSeq 4000 PE150 |
| PG_2019_TE14L01 | Red-breasted | 5/14/19 | 54.3286 | -122.6668 | u | tri | HiSeq 4000 PE150 |
| PG_2019_TE16L01a | UKSA | 5/16/19 | 53.6088 | -122.6148 | u | tri | HiSeq 4000 PE150 |
| PG_2019_TE16L01b | UKSA | 6/16/19 | 53.7394 | -122.3349 | u | tri | HiSeq 4000 PE150 |
| PG_2019_TE16L02 | Red-breasted | 5/16/19 | 53.6088 | -122.6148 | u | tri | HiSeq 4000 PE150 |
| PG_2019_TE16L03 | Red-breasted | 5/16/19 | 53.6019 | -122.5772 | u | tri | HiSeq 4000 PE150 |
| PG_2019_TE17L01 | Red-breasted | 5/17/19 | 53.7282 | -120.8787 | u | tri | HiSeq 4000 PE150 |
| PG_2019_TE17L02 | Red-breasted | 5/17/19 | 53.7282 | -120.8787 | u | tri | HiSeq 4000 PE150 |
| PG_2019_TE17L03 | Red-breasted | 5/17/19 | 53.7057 | -120.8983 | u | tri | HiSeq 4000 PE150 |
| PG_2019_TE19L01 | Red-breasted | 5/19/19 | 53.9669 | -124.3667 | u | tri | HiSeq 4000 PE150 |
| PG_2019_TE19L03 | Yellow-bellied | 5/19/19 | 53.9625 | -124.3535 | m | tri | HiSeq 4000 PE150 |
| PG_2019_TE19L04 | Yellow-bellied | 5/19/19 | 53.9625 | -124.3535 | m | tri | HiSeq 4000 PE150 |
| PG_2019_TE20L01 | Red-breasted | 5/20/19 | 53.3613 | -122.598 | u | tri | HiSeq 4000 PE150 |
| PG_2019_TE20L02 | Red-breasted | 5/20/19 | 53.3613 | -122.598 | u | tri | HiSeq 4000 PE150 |
| PG_2019_TE21L01 | Red-breasted | 5/21/19 | 53.91126 | -122.27753 | u | tri | HiSeq 4000 PE150 |
| PG_2019_TE21L02 | Hybrid | 5/21/19 | 53.8937 | -122.3466 | u | tri | HiSeq 4000 PE150 |
| PG_2019_TE23L01 | Yellow-bellied | 5/23/19 | 54.2103 | -122.5386 | m | tri | HiSeq 4000 PE150 |
| PG_2019_TE23L02 | Red-breasted | 5/23/19 | 54.0209 | -122.7351 | u | tri | HiSeq 4000 PE150 |
| PG_2019_TE23L03 | Red-breasted | 5/23/19 | 54.0209 | -122.7351 | u | tri | HiSeq 4000 PE150 |
| PG_2019_TE24L01 | Red-breasted | 5/24/19 | 54.1442 | -122.2198 | u | tri | HiSeq 4000 PE150 |
| PG_2019_TE24L02 | Red-breasted | 5/24/19 | 54.1415 | -124.2045 | u | tri | HiSeq 4000 PE150 |
| PG_2019_TE25L01 | Red-breasted | 5/25/19 | 53.7771 | -122.7003 | u | tri | HiSeq 4000 PE150 |
| PG_2019_TE25L02 | Red-breasted | 5/25/19 | 53.7716 | -122.6729 | u | tri | HiSeq 4000 PE150 |
| PG_2019_TE25L03 | Red-breasted | 5/25/19 | 53.7716 | -122.6729 | u | tri | HiSeq 4000 PE150 |
| PG_2019_TE26L01 | Red-breasted | 5/26/19 | 53.7314 | -122.2653 | u | tri | HiSeq 4000 PE150 |
| PG_2019_TE26L02 | Red-breasted | 5/26/19 | 53.7314 | -122.2653 | u | tri | HiSeq 4000 PE150 |
| PG_2019_TE27L01 | Red-breasted | 5/27/19 | 54.2415 | -122.5295 | u | tri | HiSeq 4000 PE150 |
| PG_2019_TE27L02 | Red-breasted | 5/27/19 | 54.2403 | -122.5204 | u | tri | HiSeq 4000 PE150 |
| PG_2019_TE27L03 | Hybrid | 5/27/19 | 54.2303 | -122.6021 | u | tri | HiSeq 4000 PE150 |
| PG_2019_TE28L02 | Hybrid | 5/28/19 | 53.8875 | -122.8211 | u | tri | HiSeq 4000 PE150 |
| PG_2019_TE28L03 | Red-breasted | 5/28/19 | 53.6625 | -122.9519 | u | tri | HiSeq 4000 PE150 |
| PG_2019_TE28L04 | Hybrid | 5/28/19 | 53.6407 | -122.9818 | u | tri | HiSeq 4000 PE150 |
| PG_2019_TE29L01 | Hybrid | 5/29/19 | 53.1204 | -122.3083 | u | tri | HiSeq 4000 PE150 |
| PG_2019_TE29L02 | Hybrid | 5/29/19 | 53.0449 | -122.6035 | u | tri | HiSeq 4000 PE150 |
| PG_2019_TE30L01 | Hybrid | 5/30/19 | 53.8699 | -121.9677 | u | tri | HiSeq 4000 PE150 |
| PG_2019_TE31L01 | Yellow-bellied | 5/31/19 | 54.4928 | -122.7664 | m | tri | HiSeq 4000 PE150 |
| PG_2019_TF01L01 | Hybrid | 6/1/19 | 53.8382 | -123.1933 | u | tri | HiSeq 4000 PE150 |
| PG_2019_TF03L01 | Red-breasted | 6/3/19 | 53.0891 | -122.6134 | u | tri | HiSeq 4000 PE150 |
| PG_2019_TF04L01 | Hybrid | 6/4/19 | 54.0212 | -122.2926 | u | tri | HiSeq 4000 PE150 |
| PG_2019_TF04L02 | Yellow-bellied | 6/4/19 | 54.0084 | -122.266 | m | tri | HiSeq 4000 PE150 |
| PG_2019_TF05L01 | Red-breasted | 6/5/19 | 54.5993 | -123.0668 | u | tri | HiSeq 4000 PE150 |
| PG_2019_TF05L02 | Red-breasted | 6/5/19 | 54.5993 | -123.0668 | u | tri | HiSeq 4000 PE150 |
| PG_2019_TF06L01 | Yellow-bellied | 6/6/19 | 53.0481 | -122.4572 | m | tri | HiSeq 4000 PE150 |
| PG_2019_TF09L01 | Red-breasted | 6/9/19 | 54.1817 | -122.6354 | u | tri | HiSeq 4000 PE150 |
| PG_2019_TF09L02 | Red-breasted | 6/9/19 | 54.1817 | -122.6354 | u | tri | HiSeq 4000 PE150 |
| PG_2019_TF09L03 | Red-breasted | 6/9/19 | 54.1842 | -122.6452 | u | tri | HiSeq 4000 PE150 |
| PG_2019_TF13L01 | Red-breasted | 6/13/19 | 53.5489 | -122.656 | u | tri | HiSeq 4000 PE150 |
| PG_2019_TF13L02 | Red-breasted | 6/13/19 | 53.5489 | -122.656 | u | tri | HiSeq 4000 PE150 |
| PG_2019_TF14L01 | Hybrid | 6/14/19 | 53.7551 | -123.1776 | u | tri | HiSeq 4000 PE150 |
| PG_2019_TF14L02 | Red-breasted | 6/14/19 | 53.7551 | -123.1776 | u | tri | HiSeq 4000 PE150 |
| PG_2019_TF15L01 | Red-breasted | 6/15/19 | 54.0822 | -122.3662 | u | tri | HiSeq 4000 PE150 |
| PG_2019_TF15L02 | Red-breasted | 6/15/19 | 54.0822 | -122.3662 | u | tri | HiSeq 4000 PE150 |
| PG_2019_TF15L03 | Yellow-bellied | 6/15/19 | 54.1097 | -122.3853 | f | tri | HiSeq 4000 PE150 |
| PG_2019_TF15L04 | Yellow-bellied | 6/15/19 | 54.1097 | -122.3853 | m | tri | HiSeq 4000 PE150 |
| PG_2019_TF15L05 | Yellow-bellied | 6/15/19 | 54.1097 | -122.3853 | m | tri | HiSeq 4000 PE150 |
| PG_2019_TF17L01 | Red-breasted | 6/17/19 | 53.7492 | -123.1908 | u | tri | HiSeq 4000 PE150 |
| PG_2019_TF17L02 | Red-breasted | 6/17/19 | 53.7492 | -123.1908 | u | tri | HiSeq 4000 PE150 |
| PG_2019_TF19L01 | Red-naped | 6/19/19 | 52.7817 | -122.3334 | f | tri | HiSeq 4000 PE150 |
| PG_2019_TF19L02 | Hybrid | 6/19/19 | 52.7817 | -122.3334 | m | tri | HiSeq 4000 PE150 |
| PG_2019_TF19L03 | Red-naped | 6/19/19 | 52.7639 | -122.3346 | m | tri | HiSeq 4000 PE150 |
| PG_2019_TF20L01 | Hybrid | 6/20/19 | 52.8922 | -122.5707 | u | tri | HiSeq 4000 PE150 |
| PG_2019_TF21L01 | Red-naped | 6/21/19 | 52.8331 | -122.622 | m | tri | HiSeq 4000 PE150 |
| PG_2019_TF21L02 | Red-naped | 6/21/19 | 52.8331 | -122.622 | f | tri | HiSeq 4000 PE150 |
| PG_2019_TF23L01 | Red-naped | 6/23/19 | 52.6479 | -122.4427 | m | tri | HiSeq 4000 PE150 |
| PG_2019_TF23L02 | Red-naped | 6/23/19 | 52.6443 | -122.4045 | m | tri | HiSeq 4000 PE150 |
| PG_2019_TF23L03 | Red-naped | 6/23/19 | 52.6443 | -122.4045 | m | tri | HiSeq 4000 PE150 |
| PG_2019_TF24L01 | Red-breasted | 6/24/19 | 52.9268 | -122.6182 | u | tri | HiSeq 4000 PE150 |
| PG_2019_TF25L01 | Red-breasted | 6/25/19 | 53.0185 | -122.08 | u | tri | HiSeq 4000 PE150 |
| PG_2019_TF26L01 | Hybrid | 6/26/19 | 53.8346 | -123.5957 | u | tri | HiSeq 4000 PE150 |
| PG_2019_TF26L02 | Red-breasted | 6/26/19 | 53.8346 | -123.5957 | u | tri | HiSeq 4000 PE150 |
| PG_2019_TF26L03 | Hybrid | 6/26/19 | 53.8382 | -123.1933 | u | tri | HiSeq 4000 PE150 |
| PG_2020_TE06L01 | Red-breasted | 5/6/19 | 53.8765 | -122.272 | u | tri | NovaSeqSP 6000 PE150 |
| PG_2020_TE07L03 | Hybrid | 5/7/19 | 53.76997 | -122.63411 | u | tri | NovaSeqSP 6000 PE150 |
| PG_2020_TE16L01 | Red-breasted | 5/16/19 | 53.60883 | -122.61477 | u | tri | NovaSeqSP 6000 PE150 |
| PG_2020_TE19L02 | Red-breasted | 5/19/19 | 53.9669 | -124.3667 | u | tri | NovaSeqSP 6000 PE150 |
| PG_2020_TE21L01 | Red-breasted | 5/21/19 | 53.9113 | -122.2775 | u | tri | NovaSeqSP 6000 PE150 |
| PG_2020_TE24L02 | Red-breasted | 5/24/19 | 54.141492 | -124.20448 | u | tri | NovaSeqSP 6000 PE150 |
| PG_2020_TE27L02 | Red-breasted | 5/27/19 | 54.240305 | -122.52039 | u | tri | NovaSeqSP 6000 PE150 |
| PG_2020_TE28L01 | Yellow-bellied | 5/28/19 | 53.8875 | -122.8211 | m | tri | NovaSeqSP 6000 PE150 |
| PG_2020_TE29L01 | Hybrid | 5/29/19 | 53.120402 | -122.30834 | u | tri | NovaSeqSP 6000 PE150 |
| PG_2020_TE31L01 | Yellow-bellied | 5/31/19 | 54.492837 | -122.76638 | m | tri | NovaSeqSP 6000 PE150 |
| PG_2020_TF10L01 | Red-breasted | 6/10/19 | 54.1998 | -122.694 | u | tri | NovaSeqSP 6000 PE150 |
| PG_2020_TF21L01 | Red-naped | 6/21/19 | 52.83305 | -122.62201 | m | tri | NovaSeqSP 6000 PE150 |
| PG_2020_TF23L04_ff | Hybrid | 6/23/19 | 52.6346 | -122.3997 | m | tri | NovaSeqSP 6000 PE150 |
| PG_2020_TF26L03 | Hybrid | 6/26/19 | 53.8382 | -123.19329 | u | tri | NovaSeqSP 6000 PE150 |
| PG_2020_UE12L01 | Red-breasted | 5/12/20 | 53.2283 | -122.4202 | u | tri | NovaSeqSP 6000 PE150 |
| PG_2020_UE12L02 | Red-breasted | 5/12/20 | 53.2283 | -122.4202 | u | tri | NovaSeqSP 6000 PE150 |
| PG_2020_UE13L01 | Hybrid | 5/13/20 | 53.3006 | -122.4331 | m | tri | NovaSeqSP 6000 PE150 |
| PG_2020_UE14L01 | Hybrid | 5/14/20 | 53.4482 | -122.659 | u | tri | NovaSeqSP 6000 PE150 |
| PG_2020_UE14L02 | Hybrid | 5/14/20 | 53.4722 | -122.5819 | u | tri | NovaSeqSP 6000 PE150 |
| PG_2020_UE14L03 | Red-breasted | 5/14/20 | 53.2812 | -122.5695 | u | tri | NovaSeqSP 6000 PE150 |
| PG_2020_UE14L04 | Red-breasted | 5/14/20 | 53.2812 | -122.5695 | u | tri | NovaSeqSP 6000 PE150 |
| PG_2020_UE15L01 | Red-breasted | 5/15/20 | 53.1675 | -122.4895 | f | tri | NovaSeqSP 6000 PE150 |
| PG_2020_UE15L02 | Red-breasted | 5/15/20 | 53.1675 | -122.4895 | u | tri | NovaSeqSP 6000 PE150 |
| PG_2020_UE16L01 | Red-breasted | 5/16/20 | 53.8649 | -121.5251 | u | tri | NovaSeqSP 6000 PE150 |
| PG_2020_UE17L01 | Hybrid | 5/17/20 | 53.4014 | -120.2892 | u | tri | NovaSeqSP 6000 PE150 |
| PG_2020_UE17L02 | Hybrid | 5/17/20 | 53.4048 | -120.3025 | u | tri | NovaSeqSP 6000 PE150 |
| PG_2020_UE18L01 | Red-breasted | 5/18/20 | 53.6165 | -120.6701 | u | tri | NovaSeqSP 6000 PE150 |
| PG_2020_UE19L01 | Hybrid | 5/19/20 | 53.8337 | -121.3357 | f | tri | NovaSeqSP 6000 PE150 |
| PG_2020_UE20L01 | Red-breasted | 5/20/20 | 53.8362 | -121.7826 | u | tri | NovaSeqSP 6000 PE150 |
| woodpeckers_SS_1E27S01 | Yellow-bellied | 5/27/09 | 55.3911 | -123.1786 | m | di-RBxYB hybrid | HiSeq 2000 PE150 |
| woodpeckers_SS_1E27S03 | Yellow-bellied | 5/27/09 | 55.3911 | -123.1786 | m | di-RBxYB hybrid | HiSeq 2000 PE150 |
| woodpeckers_SS_1E27S04 | Yellow-bellied | 5/27/09 | 55.3911 | -123.1786 | m | di-RBxYB hybrid | HiSeq 2000 PE150 |
| woodpeckers_SS_1E27S06 | Yellow-bellied | 5/27/09 | 55.4675 | -123.2414 | m | di-RBxYB hybrid | HiSeq 2000 PE150 |
| woodpeckers_SS_freez3 | Yellow-bellied | NA | 53.5333 | -113.0166 | f | allo-yb | HiSeq 2000 PE150 |
| woodpeckers_SS_freez4 | Yellow-bellied | NA | 53.89 | -113.67 | f | allo-yb | HiSeq 2000 PE150 |
| woodpeckers_SS_freez5 | Yellow-bellied | NA | 53.53 | -113.51 | m | allo-yb | HiSeq 2000 PE150 |
| woodpeckers_SS_freez7 | Yellow-bellied | NA | 53.41 | -112.9 | f | allo-yb | HiSeq 2000 PE150 |
| woodpeckers_SS_IE08S02 | Yellow-bellied | 5/8/09 | 55.7411 | -121.8353 | m | di-RBxYB hybrid | HiSeq 2000 PE150 |
| woodpeckers_SS_IE09S01 | Yellow-bellied | 5/9/09 | 55.8867 | -121.6586 | m | di-RBxYB hybrid | HiSeq 2000 PE150 |
| woodpeckers_SS_IE09S02 | Yellow-bellied | 5/9/09 | 55.8186 | -121.6886 | m | di-RBxYB hybrid | HiSeq 2000 PE150 |
| woodpeckers_SS_IE10S01 | Yellow-bellied | 5/10/09 | 55.9039 | -121.6481 | m | di-RBxYB hybrid | HiSeq 2000 PE150 |
| woodpeckers_SS_IE10S02 | Yellow-bellied | 5/10/09 | 55.8794 | -121.6169 | m | di-RBxYB hybrid | HiSeq 2000 PE150 |
| woodpeckers_SS_IE10S03 | Yellow-bellied | 5/10/09 | 55.7225 | -121.5633 | m | di-RBxYB hybrid | HiSeq 2000 PE150 |
| woodpeckers_SS_IE10S04 | Yellow-bellied | 5/10/09 | 55.6969 | -121.73 | f | di-RBxYB hybrid | HiSeq 2000 PE150 |
| woodpeckers_SS_IE11S04 | Yellow-bellied | 5/11/09 | 55.705 | -121.7367 | m | di-RBxYB hybrid | HiSeq 2000 PE150 |
| woodpeckers_SS_IE14S01 | Yellow-bellied | 5/14/09 | 55.8686 | -122.2872 | m | di-RBxYB hybrid | HiSeq 2000 PE150 |
| woodpeckers_SS_IE14S02 | Yellow-bellied | 5/14/09 | 55.7217 | -122.4264 | m | di-RBxYB hybrid | HiSeq 2000 PE150 |
| woodpeckers_SS_IE15S01 | Yellow-bellied | 5/15/09 | 55.6508 | -122.3758 | m | di-RBxYB hybrid | HiSeq 2000 PE150 |
| woodpeckers_SS_IE15S03 | Yellow-bellied | 5/15/09 | 55.7503 | -122.4786 | m | di-RBxYB hybrid | HiSeq 2000 PE150 |
| woodpeckers_SS_IE16S01 | Red-breasted | 5/16/09 | 55.6536 | -122.5453 | u | di-RBxYB hybrid | HiSeq 2000 PE150 |
| woodpeckers_SS_IE18S01 | Yellow-bellied | 5/18/09 | 55.8267 | -122.2767 | m | di-RBxYB hybrid | HiSeq 2000 PE150 |
| woodpeckers_SS_IE18S04 | Yellow-bellied | 5/18/09 | 55.875 | -122.4514 | m | di-RBxYB hybrid | HiSeq 2000 PE150 |
| woodpeckers_SS_IE20S01 | Yellow-bellied | 5/20/09 | 55.7664 | -122.3325 | m | di-RBxYB hybrid | HiSeq 2000 PE150 |
| woodpeckers_SS_IE20S02 | Yellow-bellied | 5/20/09 | 55.7072 | -122.5142 | m | di-RBxYB hybrid | HiSeq 2000 PE150 |
| woodpeckers_SS_IE20S03 | Yellow-bellied | 5/20/09 | 55.6197 | -122.5167 | m | di-RBxYB hybrid | HiSeq 2000 PE150 |
| woodpeckers_SS_IE20S04 | Yellow-bellied | 5/20/09 | 55.69 | -122.4972 | f | di-RBxYB hybrid | HiSeq 2000 PE150 |
| woodpeckers_SS_IE21S01 | Yellow-bellied | 5/21/09 | 55.7117 | -122.5075 | m | di-RBxYB hybrid | HiSeq 2000 PE150 |
| woodpeckers_SS_IE21S02 | Yellow-bellied | 5/21/09 | 55.7117 | -122.5075 | f | di-RBxYB hybrid | HiSeq 2000 PE150 |
| woodpeckers_SS_IE24S01 | Hybrid | 5/22/09 | 55.4953 | -123.1794 | u | di-RBxYB hybrid | HiSeq 2000 PE150 |
| woodpeckers_SS_IE24S02 | Yellow-bellied | 5/23/09 | 54.01 | -122.9247 | m | di-RBxYB hybrid | HiSeq 2000 PE150 |
| woodpeckers_SS_IE24S03 | RBxYB hybrid | 5/24/09 | 55.4953 | -123.1794 | m | di-RBxYB hybrid | HiSeq 2000 PE150 |
| woodpeckers_SS_IE25S01 | Yellow-bellied | 5/24/09 | 55.5889 | -123.12 | m | di-RBxYB hybrid | HiSeq 2000 PE150 |
| woodpeckers_SS_IE25S02 | Yellow-bellied | 5/24/09 | 55.4389 | -123.2233 | m | di-RBxYB hybrid | HiSeq 2000 PE150 |
| woodpeckers_SS_IE25S03 | Yellow-bellied | 5/25/09 | 55.6214 | -123.2083 | m | di-RBxYB hybrid | HiSeq 2000 PE150 |
| woodpeckers_SS_IE28S02 | Yellow-bellied | 5/27/09 | 55.4675 | -123.2414 | m | di-RBxYB hybrid | HiSeq 2000 PE150 |
| woodpeckers_SS_IE28S03 | Red-breasted | 5/28/09 | 55.3319 | -123.4075 | f | di-RBxYB hybrid | HiSeq 2000 PE150 |
| woodpeckers_SS_IF01S02 | Red-breasted | 5/30/09 | 55.2547 | -123.2639 | f | di-RBxYB hybrid | HiSeq 2000 PE150 |
| woodpeckers_SS_IF02S02 | Red-breasted | 6/1/09 | 55.1225 | -123.2722 | m | di-RBxYB hybrid | HiSeq 2000 PE150 |
| woodpeckers_SS_IF03S01 | RBxYB hybrid | 6/2/09 | 55.7008 | -123.3733 | m | di-RBxYB hybrid | HiSeq 2000 PE150 |
| woodpeckers_SS_IF03S02 | Red-breasted | 6/2/09 | 55.6442 | -123.3808 | m | di-RBxYB hybrid | HiSeq 2000 PE150 |
| woodpeckers_SS_IF03S03 | Yellow-bellied | 6/3/09 | 55.2686 | -123.3972 | f | di-RBxYB hybrid | HiSeq 2000 PE150 |
| woodpeckers_SS_IF03S04 | RBxYB hybrid | 6/3/09 | 55.2686 | -123.3972 | m | di-RBxYB hybrid | HiSeq 2000 PE150 |
| woodpeckers_SS_IF05S01 | Red-breasted | 6/3/09 | 55.4611 | -123.36 | m | di-RBxYB hybrid | HiSeq 2000 PE150 |
| woodpeckers_SS_IF05S02 | Red-breasted | 6/3/09 | 55.4611 | -123.36 | f | di-RBxYB hybrid | HiSeq 2000 PE150 |
| woodpeckers_SS_IF05S03 | Red-breasted | 6/5/09 | 54.8494 | -124.5533 | f | di-RBxYB hybrid | HiSeq 2000 PE150 |
| woodpeckers_SS_IF05S04 | Red-breasted | 6/5/09 | 54.8494 | -124.5533 | m | di-RBxYB hybrid | HiSeq 2000 PE150 |
| woodpeckers_SS_IF06S01 | Red-breasted | 6/5/09 | 54.8806 | -124.6044 | m | di-RBxYB hybrid | HiSeq 2000 PE150 |
| woodpeckers_SS_IF06S02 | Red-breasted | 6/5/09 | 54.7117 | -124.4614 | u | di-RBxYB hybrid | HiSeq 2000 PE150 |
| woodpeckers_SS_IF06S03 | Red-breasted | 6/6/09 | 54.685 | -124.6125 | m | di-RBxYB hybrid | HiSeq 2000 PE150 |
| woodpeckers_SS_IF08S01 | Red-breasted | 6/6/09 | 54.8333 | -124.6164 | m | di-RBxYB hybrid | HiSeq 2000 PE150 |
| woodpeckers_SS_IF08S03 | Red-breasted | 6/8/09 | 54.8297 | -124.3103 | u | di-RBxYB hybrid | HiSeq 2000 PE150 |
| woodpeckers_SS_IF08S04 | Yellow-bellied | 6/8/09 | 54.8297 | -124.3103 | m | di-RBxYB hybrid | HiSeq 2000 PE150 |
| woodpeckers_SS_IF08S05 | Red-breasted | 6/8/09 | 54.5983 | -124.1542 | m | di-RBxYB hybrid | HiSeq 2000 PE150 |
| woodpeckers_SS_IF09S01 | Red-breasted | 6/8/09 | 54.6731 | -124.1892 | f | di-RBxYB hybrid | HiSeq 2000 PE150 |
| woodpeckers_SS_IF10S02 | Red-breasted | 6/9/09 | 54.575 | -124.4922 | m | di-RBxYB hybrid | HiSeq 2000 PE150 |
| woodpeckers_SS_IF11S01 | Red-breasted | 6/10/09 | 54.4194 | -124.3589 | u | di-RBxYB hybrid | HiSeq 2000 PE150 |
| woodpeckers_SS_IF12S03 | Red-breasted | 6/12/09 | 54.8558 | -124.1906 | m | di-RBxYB hybrid | HiSeq 2000 PE150 |
| woodpeckers_SS_IF12S04 | Red-breasted | 6/12/09 | 54.8558 | -124.1906 | f | di-RBxYB hybrid | HiSeq 2000 PE150 |
| woodpeckers_SS_IF12S05 | Red-breasted | 6/12/09 | 54.9031 | -124.3308 | u | di-RBxYB hybrid | HiSeq 2000 PE150 |
| woodpeckers_SS_IF12S06 | Red-breasted | 6/12/09 | 55.0181 | -124.1628 | m | di-RBxYB hybrid | HiSeq 2000 PE150 |
| woodpeckers_SS_IF13S01 | Red-breasted | 6/13/09 | 55.0061 | -124.1892 | u | di-RBxYB hybrid | HiSeq 2000 PE150 |
| woodpeckers_SS_IF13S02 | Yellow-bellied | 6/13/09 | 55.0061 | -124.1892 | f | di-RBxYB hybrid | HiSeq 2000 PE150 |
| woodpeckers_SS_IF13S03 | Red-breasted | 6/13/09 | 54.915 | -124.195 | u | di-RBxYB hybrid | HiSeq 2000 PE150 |
| woodpeckers_SS_IF16S02 | Red-breasted | 6/16/09 | 54.1503 | -126.6156 | m | di-RBxYB hybrid | HiSeq 2000 PE150 |
| woodpeckers_SS_IF16S03 | Red-breasted | 6/16/09 | 54.1275 | -126.7197 | m | di-RBxYB hybrid | HiSeq 2000 PE150 |
| woodpeckers_SS_IF16S04 | Red-breasted | 6/16/09 | 54.0481 | -126.6503 | u | di-RBxYB hybrid | HiSeq 2000 PE150 |
| woodpeckers_SS_IF16S05 | Red-breasted | 6/16/09 | 54.0481 | -126.6503 | u | di-RBxYB hybrid | HiSeq 2000 PE150 |
| woodpeckers_SS_IF17S01 | Red-breasted | 6/17/09 | 54.0564 | -126.6594 | f | di-RBxYB hybrid | HiSeq 2000 PE150 |
| woodpeckers_SS_IF17S02 | Red-breasted | 6/17/09 | 54.0564 | -126.6594 | m | di-RBxYB hybrid | HiSeq 2000 PE150 |
| woodpeckers_SS_JF13S02 | RNxRB hybrid | 6/13/10 | 52.2153 | -122.4047 | m | di-RNxRB hybrid | HiSeq 2000 PE150 |
| woodpeckers_SS_JF13S03 | Red-naped | 6/13/10 | 52.1514 | -122.6058 | m | di-RNxRB hybrid | HiSeq 2000 PE150 |
| woodpeckers_SS_JF14S01 | RNxRB hybrid | 6/14/10 | 52.1469 | -122.5419 | m | di-RNxRB hybrid | HiSeq 2000 PE150 |
| woodpeckers_SS_JF14S02 | Red-naped | 6/14/10 | 52.1469 | -122.5419 | f | di-RNxRB hybrid | HiSeq 2000 PE150 |
| woodpeckers_SS_JF15S01 | Red-naped | 6/15/10 | 52.1319 | -122.5739 | f | di-RNxRB hybrid | HiSeq 2000 PE150 |
| woodpeckers_SS_JF15S02 | RNxRB hybrid | 6/15/10 | 52.1319 | -122.5739 | m | di-RNxRB hybrid | HiSeq 2000 PE150 |
| woodpeckers_SS_JF16S02 | RNxRB hybrid | 6/16/10 | 52.2792 | -121.9317 | m | di-RNxRB hybrid | HiSeq 2000 PE150 |
| woodpeckers_SS_JF16S03 | Red-naped | 6/16/10 | 52.2792 | -121.9317 | f | di-RNxRB hybrid | HiSeq 2000 PE150 |
| woodpeckers_SS_JF16S04 | Red-naped | 6/16/10 | 52.4289 | -122.1 | f | di-RNxRB hybrid | HiSeq 2000 PE150 |
| woodpeckers_SS_JF17S01 | RNxRB hybrid | 6/17/10 | 52.0814 | -122.6103 | m | di-RNxRB hybrid | HiSeq 2000 PE150 |
| woodpeckers_SS_JF17S02 | RNxRB hybrid | 6/17/10 | 52.0814 | -122.6103 | f | di-RNxRB hybrid | HiSeq 2000 PE150 |
| woodpeckers_SS_JF17S06 | RNxRB hybrid | 6/17/10 | 52.0361 | -122.4097 | m | di-RNxRB hybrid | HiSeq 2000 PE150 |
| woodpeckers_SS_JL01S01 | Red-breasted | 12/1/10 | 49.3043 | -123.1443 | u | allo-rb | HiSeq 2000 PE150 |
| woodpeckers_SS_KE18S01 | RNxRB hybrid | 5/18/11 | 52.2231 | -125.1122 | m | di-RNxRB hybrid | HiSeq 2000 PE150 |
| woodpeckers_SS_KE18S02 | RNxRB hybrid | 5/18/11 | 52.2206 | -124.9242 | m | di-RNxRB hybrid | HiSeq 2000 PE150 |
| woodpeckers_SS_KE18S03 | Red-naped | 5/18/11 | 52.1608 | -125.0078 | m | di-RNxRB hybrid | HiSeq 2000 PE150 |
| woodpeckers_SS_KE18S04 | Red-naped | 5/18/11 | 52.1608 | -125.0078 | f | di-RNxRB hybrid | HiSeq 2000 PE150 |
| woodpeckers_SS_KE19S01 | Red-naped | 5/19/11 | 52.1753 | -125.1839 | m | di-RNxRB hybrid | HiSeq 2000 PE150 |
| woodpeckers_SS_KE19S02 | RNxRB hybrid | 5/19/11 | 52.1753 | -125.1839 | f | di-RNxRB hybrid | HiSeq 2000 PE150 |
| woodpeckers_SS_KE20S01 | Red-breasted | 5/20/11 | 52.8581 | -125.5617 | m | di-RNxRB hybrid | HiSeq 2000 PE150 |
| woodpeckers_SS_KE20S02 | Red-breasted | 5/20/11 | 52.8581 | -125.5617 | f | di-RNxRB hybrid | HiSeq 2000 PE150 |
| woodpeckers_SS_KE21S01 | Red-breasted | 5/21/11 | 52.5897 | -126.275 | m | di-RNxRB hybrid | HiSeq 2000 PE150 |
| woodpeckers_SS_KE21S02 | Red-breasted | 5/21/11 | 52.4167 | -126.1389 | m | di-RNxRB hybrid | HiSeq 2000 PE150 |
| woodpeckers_SS_KE22S01 | Red-breasted | 5/22/11 | 52.5933 | -126.0089 | m | di-RNxRB hybrid | HiSeq 2000 PE150 |
| woodpeckers_SS_KE22S02 | Red-breasted | 5/22/11 | 52.5425 | -126.1586 | m | di-RNxRB hybrid | HiSeq 2000 PE150 |
| woodpeckers_SS_KE22S03 | Red-breasted | 5/22/11 | 52.6206 | -125.9892 | m | di-RNxRB hybrid | HiSeq 2000 PE150 |
| woodpeckers_SS_KE22S04 | Red-breasted | 5/22/11 | 52.5375 | -126.0894 | m | di-RNxRB hybrid | HiSeq 2000 PE150 |
| woodpeckers_SS_KE24S01 | Red-breasted | 5/24/11 | 52.5331 | -126.5353 | m | di-RNxRB hybrid | HiSeq 2000 PE150 |
| woodpeckers_SS_KE25S01 | Red-breasted | 5/25/11 | 52.4533 | -126.8839 | m | di-RNxRB hybrid | HiSeq 2000 PE150 |
| woodpeckers_SS_KE25S03 | Red-breasted | 5/25/11 | 52.4903 | -126.9036 | m | di-RNxRB hybrid | HiSeq 2000 PE150 |
| woodpeckers_SS_KE27S01 | Red-breasted | 5/27/11 | 52.535 | -126.7636 | f | di-RNxRB hybrid | HiSeq 2000 PE150 |
| woodpeckers_SS_KE27S02 | Red-breasted | 5/27/11 | 52.6217 | -126.8753 | f | di-RNxRB hybrid | HiSeq 2000 PE150 |
| woodpeckers_SS_KE27S04 | Red-breasted | 5/27/11 | 52.65 | -126.6394 | m | di-RNxRB hybrid | HiSeq 2000 PE150 |
| woodpeckers_SS_KE29S01 | Red-breasted | 5/29/11 | 52.4808 | -126.9217 | m | di-RNxRB hybrid | HiSeq 2000 PE150 |
| woodpeckers_SS_KE29S02 | Red-breasted | 5/29/11 | 52.4808 | -126.9217 | f | di-RNxRB hybrid | HiSeq 2000 PE150 |
| woodpeckers_SS_KE29S04 | Red-breasted | 5/29/11 | 52.4667 | -126.8658 | m | di-RNxRB hybrid | HiSeq 2000 PE150 |
| woodpeckers_SS_KE29S05 | Red-breasted | 5/29/11 | 52.5328 | -126.4631 | m | di-RNxRB hybrid | HiSeq 2000 PE150 |
| woodpeckers_SS_KE29S06 | Red-breasted | 5/29/11 | 52.5328 | -126.4631 | f | di-RNxRB hybrid | HiSeq 2000 PE150 |
| woodpeckers_SS_KE30S01 | Red-breasted | 5/30/11 | 52.6042 | -126.7286 | m | di-RNxRB hybrid | HiSeq 2000 PE150 |
| woodpeckers_SS_KE30S02 | Red-breasted | 5/30/11 | 52.6042 | -126.7286 | f | di-RNxRB hybrid | HiSeq 2000 PE150 |
| woodpeckers_SS_KE31S02 | RNxRB hybrid | 5/31/11 | 51.9172 | -124.7106 | m | di-RNxRB hybrid | HiSeq 2000 PE150 |
| woodpeckers_SS_KF03S01 | Red-naped | 6/3/11 | 51.8658 | -121.4275 | m | di-RNxRB hybrid | HiSeq 2000 PE150 |
| woodpeckers_SS_KF04S01 | Red-naped | 6/4/11 | 51.7361 | -121.1144 | m | di-RNxRB hybrid | HiSeq 2000 PE150 |
| woodpeckers_SS_KF04S07 | Red-naped | 6/4/11 | 51.6169 | -120.4364 | m | di-RNxRB hybrid | HiSeq 2000 PE150 |
| woodpeckers_SS_KF05S01 | Red-naped | 6/5/11 | 51.6936 | -120.4111 | m | di-RNxRB hybrid | HiSeq 2000 PE150 |
| woodpeckers_SS_KF06S01 | Red-naped | 6/6/11 | 50.5103 | -117.4319 | m | allo-rn | HiSeq 2000 PE150 |
| woodpeckers_SS_KF06S02 | Red-naped | 6/6/11 | 50.5569 | -117.3556 | m | allo-rn | HiSeq 2000 PE150 |
| woodpeckers_SS_KF06S03 | Red-naped | 6/6/11 | 50.6036 | -117.3917 | m | allo-rn | HiSeq 2000 PE150 |
| woodpeckers_SS_KF06S04 | Red-naped | 6/6/11 | 50.6036 | -117.3917 | m | allo-rn | HiSeq 2000 PE150 |
| woodpeckers_SS_KF07S01 | Red-naped | 6/7/11 | 50.3497 | -116.9978 | m | allo-rn | HiSeq 2000 PE150 |
| woodpeckers_SS_KF07S02 | Red-naped | 6/7/11 | 50.2753 | -117.1086 | m | allo-rn | HiSeq 2000 PE150 |
| woodpeckers_SS_KF07S03 | Red-naped | 6/7/11 | 50.2753 | -117.1086 | m | allo-rn | HiSeq 2000 PE150 |
| woodpeckers_SS_KF09S01 | Red-naped | 6/9/11 | 49.5644 | -117.3436 | m | allo-rn | HiSeq 2000 PE150 |
| woodpeckers_SS_KF09S02 | Red-naped | 6/9/11 | 49.2594 | -117.4214 | m | allo-rn | HiSeq 2000 PE150 |
| woodpeckers_SS_KF09S03 | Red-naped | 6/9/11 | 49.3158 | -117.4947 | m | allo-rn | HiSeq 2000 PE150 |
| woodpeckers_SS_KF09S04 | Red-naped | 6/9/11 | 49.2478 | -117.4008 | m | allo-rn | HiSeq 2000 PE150 |
| woodpeckers_SS_KF10S01 | Red-naped | 6/10/11 | 49.6467 | -117.5558 | m | allo-rn | HiSeq 2000 PE150 |
| woodpeckers_SS_KF10S02 | Red-naped | 6/10/11 | 49.6467 | -117.5558 | f | allo-rn | HiSeq 2000 PE150 |
| woodpeckers_SS_KF11S01 | Red-naped | 6/11/11 | 50.6364 | -119.455 | m | di-RNxRB hybrid | HiSeq 2000 PE150 |
| woodpeckers_SS_KF11S02 | Red-naped | 6/11/11 | 50.6364 | -119.455 | f | di-RNxRB hybrid | HiSeq 2000 PE150 |
| woodpeckers_SS_KF12S03 | Red-naped | 6/12/11 | 52.0464 | -123.5392 | f | di-RNxRB hybrid | HiSeq 2000 PE150 |
| woodpeckers_SS_KF12S04 | Red-naped | 6/12/11 | 52.0464 | -123.5392 | m | di-RNxRB hybrid | HiSeq 2000 PE150 |
| woodpeckers_SS_KF12S05 | Red-naped | 6/12/11 | 52.2942 | -124.3092 | f | di-RNxRB hybrid | HiSeq 2000 PE150 |
| woodpeckers_SS_KF12S06 | Red-naped | 6/12/11 | 52.5875 | -125.4767 | f | di-RNxRB hybrid | HiSeq 2000 PE150 |
| woodpeckers_SS_KF13S01 | RNxRB hybrid | 6/13/11 | 52.7661 | -125.6253 | f | di-RNxRB hybrid | HiSeq 2000 PE150 |
| woodpeckers_SS_KF13S02 | RNxRB hybrid | 6/13/11 | 52.7661 | -125.6253 | m | di-RNxRB hybrid | HiSeq 2000 PE150 |
| woodpeckers_SS_KF13S03 | Red-breasted | 6/13/11 | 52.5675 | -125.47 | u | di-RNxRB hybrid | HiSeq 2000 PE150 |
| woodpeckers_SS_KF14S01 | Red-naped | 6/14/11 | 52.865 | -125.56 | m | di-RNxRB hybrid | HiSeq 2000 PE150 |
| woodpeckers_SS_KF14S03 | Red-breasted | 6/14/11 | 52.7853 | -125.7175 | f | di-RNxRB hybrid | HiSeq 2000 PE150 |
| woodpeckers_SS_KF15S01 | Red-naped | 6/15/11 | 52.67 | -125.6511 | m | di-RNxRB hybrid | HiSeq 2000 PE150 |
| woodpeckers_SS_KF15S03 | Red-naped | 6/15/11 | 52.67 | -125.6511 | m | di-RNxRB hybrid | HiSeq 2000 PE150 |
| woodpeckers_SS_KF15S04 | Red-naped | 6/15/11 | 52.8236 | -125.5842 | m | di-RNxRB hybrid | HiSeq 2000 PE150 |
| woodpeckers_SS_KF15S05 | Red-naped | 6/15/11 | 52.8236 | -125.5842 | f | di-RNxRB hybrid | HiSeq 2000 PE150 |
| woodpeckers_SS_KF16S01 | Red-naped | 6/16/11 | 52.8767 | -125.4672 | f | di-RNxRB hybrid | HiSeq 2000 PE150 |
| woodpeckers_SS_KF16S03 | Red-naped | 6/16/11 | 52.7436 | -125.5517 | m | di-RNxRB hybrid | HiSeq 2000 PE150 |
| woodpeckers_SS_KF16S05 | Red-naped | 6/16/11 | 52.7342 | -125.5781 | m | di-RNxRB hybrid | HiSeq 2000 PE150 |
| woodpeckers_SS_KF16S06 | RNxRB hybrid | 6/16/11 | 52.7578 | -125.3867 | m | di-RNxRB hybrid | HiSeq 2000 PE150 |
| woodpeckers_SS_KF18S01 | Red-naped | 6/18/11 | 52.8 | -125.4344 | m | di-RNxRB hybrid | HiSeq 2000 PE150 |
| woodpeckers_SS_KF19S01 | Red-naped | 6/19/11 | 52.7039 | -125.5683 | f | di-RNxRB hybrid | HiSeq 2000 PE150 |
| woodpeckers_SS_KF19S02 | RNxRB hybrid | 6/19/11 | 52.6692 | -125.4758 | m | di-RNxRB hybrid | HiSeq 2000 PE150 |
| woodpeckers_SS_KF19S03 | Red-naped | 6/19/11 | 52.5006 | -125.3208 | m | di-RNxRB hybrid | HiSeq 2000 PE150 |
| woodpeckers_SS_KF19S05 | Red-naped | 6/19/11 | 52.6317 | -125.36 | m | di-RNxRB hybrid | HiSeq 2000 PE150 |
| woodpeckers_SS_KF20S01 | Red-naped | 6/20/11 | 52.5583 | -125.4411 | m | di-RNxRB hybrid | HiSeq 2000 PE150 |
| woodpeckers_SS_KF20S02 | Red-naped | 6/20/11 | 52.7067 | -125.6239 | m | di-RNxRB hybrid | HiSeq 2000 PE150 |
| woodpeckers_SS_KF20S03 | Red-naped | 6/20/11 | 52.7589 | -125.4972 | m | di-RNxRB hybrid | HiSeq 2000 PE150 |
| woodpeckers_SS_KF20S04 | Red-naped | 6/20/11 | 52.6997 | -125.55 | m | di-RNxRB hybrid | HiSeq 2000 PE150 |
| woodpeckers_SS_KF20S05 | Red-naped | 6/20/11 | 52.5617 | -125.4239 | m | di-RNxRB hybrid | HiSeq 2000 PE150 |
| woodpeckers_SS_KF20S06 | Red-naped | 6/20/11 | 52.5617 | -125.4239 | f | di-RNxRB hybrid | HiSeq 2000 PE150 |
| woodpeckers_SS_KF20S07 | RNxRB hybrid | 6/20/11 | 52.6594 | -125.3744 | u | di-RNxRB hybrid | HiSeq 2000 PE150 |
| woodpeckers_SS_KF21S01 | Red-naped | 6/21/11 | 52.5269 | -125.3339 | m | di-RNxRB hybrid | HiSeq 2000 PE150 |
| woodpeckers_SS_KF21S02 | Red-naped | 6/21/11 | 52.5269 | -125.3339 | f | di-RNxRB hybrid | HiSeq 2000 PE150 |
| woodpeckers_SS_KF21S03 | Red-breasted | 6/21/11 | 52.5617 | -125.2556 | m | di-RNxRB hybrid | HiSeq 2000 PE150 |
| woodpeckers_SS_KF21S04 | Red-naped | 6/21/11 | 52.5617 | -125.2556 | f | di-RNxRB hybrid | HiSeq 2000 PE150 |
| woodpeckers_SS_KF21S05 | Red-naped | 6/21/11 | 52.5236 | -125.3044 | m | di-RNxRB hybrid | HiSeq 2000 PE150 |
| woodpeckers_SS_KF22S01 | Red-breasted | 6/22/11 | 52.4589 | -125.3067 | m | di-RNxRB hybrid | HiSeq 2000 PE150 |
| woodpeckers_SS_RbSa05 | Red-breasted | NA | 49.3286 | -123.1602 | f | allo-rb | HiSeq 2000 PE150 |
| woodpeckers_SS_RbSa06 | Red-breasted | NA | 49.2827 | -123.1207 | f | allo-rb | HiSeq 2000 PE150 |
| woodpeckers_SS_RbSa07 | Red-breasted | NA | 49.2827 | -123.1207 | m | allo-rb | HiSeq 2000 PE150 |
| woodpeckers_SS_RbSa08 | Red-breasted | NA | 49.3184 | -122.8936 | u | allo-rb | HiSeq 2000 PE150 |
| woodpeckers_SS_z00_31_1 | Yellow-bellied | NA | 53.55 | -113.0667 | u | allo-yb | HiSeq 2000 PE150 |
| woodpeckers_SS_z09_11_2 | Yellow-bellied | NA | 53.35 | -113.1 | u | allo-yb | HiSeq 2000 PE150 |
| woodpeckers_SS_z99_14_21 | Yellow-bellied | NA | 54.1497 | -115.0331 | u | allo-yb | HiSeq 2000 PE150 |
